# Antibiotic-tolerant persisters are pervasive among clinical *Streptococcus pneumoniae* isolates and show strong condition-dependence

**DOI:** 10.1101/2022.07.15.500022

**Authors:** Nele Geerts, Linda De Vooght, Ioannis Passaris, Bram Van den Bergh, Paul Cos

## Abstract

*Streptococcus pneumoniae* is an important human pathogen, being one of the most common causes of community-acquired pneumonia and otitis media. Antibiotic resistance in *S. pneumoniae* is an emerging problem as it depletes our arsenal of effective drugs. In addition, persistence also contributes to the antibiotic crisis in many other pathogens, yet, in *S. pneumonia*e nothing is known about antibiotic-tolerant persisters. Persister cells are phenotypic variants that exist as a subpopulation within a clonal culture. Being tolerant to lethal antibiotics, they underly the chronic nature of a variety of infections and even help in acquiring genetic resistance. Here, we set out to identify and characterize persistence in *S. pneumoniae*. Specifically, we followed different strategies to overcome the self-limiting nature of *S. pneumoniae* as confounding factor in the prolonged monitoring of antibiotic survival needed to study persistence. In optimized conditions, we identified genuine persisters in various growth phases and for four relevant antibiotics through biphasic survival dynamics and heritability assays. Finally, we detected a high variety in antibiotic survival levels across a diverse collection of *S. pneumoniae* clinical isolates, which shows that a high natural diversity in persistence is widely present in *S. pneumoniae*. Collectively, this proof-of-concept significantly progresses the understanding of the importance of antibiotic persistence in *S. pneumoniae* infections which will set stage for characterizing its relevance to clinical outcomes and advocates for increased attention to the phenotype in both fundamental and clinical research.

**IMPORTANCE:** *S. pneumoniae* is considered a serious threat by the Centers of Disease Control and Prevention through arising antibiotic resistance. In addition to resistance, bacteria can also survive lethal antibiotic treatment by developing antibiotic tolerance and more specifically by antibiotic tolerance through persistence. This phenotypic variation seems omnipresent among bacterial life, is linked to therapy failure and acts as a catalyst for resistance development. This study gives the first proof of the presence of persister cells in *S. pneumoniae* and shows a high variety in persistence levels among diverse strains, suggesting persistence is a general trait in *S. pneumoniae* cultures and that a broad range of genetic elements are controlling the phenotype. Together, our work advocates for higher interest for persistence in *S. pneumoniae* as a contributing factor for therapy failure and resistance development.

## Introduction

*Streptococcus pneumoniae* is an important human pathogen causing infections of the local mucosa, like otitis media and sinusitis, and even more severe diseases like community-acquired pneumonia (CAP) and meningitis (1, 2). Yearly, 2 million people in the United States suffer from pneumococcal-related infections, resulting in $4 billion in costs (3). It is estimated that 900,000 of these infections are caused by drug-resistant strains (3). The Centers for Disease Control and Prevention considers *S. pneumoniae* a serious threat (3). In addition to resistance, bacteria can also survive lethal antibiotic treatment by developing antibiotic tolerance or the ability to survive antibiotic treatments longer, for example due to a lower killing rate without a change in the minimum inhibitory concentration (MIC) (4, 5). Despite frequently overlooked, antibiotic tolerance can set stage for the development of genetic resistance and is associated with therapy failure (5–7). Regardless of numerous reports of persistence in a variety of bacterial species (8–11), very little is known about antibiotic tolerance, and, to the best of our knowledge, nothing is known about antibiotic tolerance through persistence, in *S. pneumoniae* (12–16).

Persistent bacteria are a subpopulation of cells that transiently switch to a non-growing state that enables them to survive treatment with a bactericidal drug concentration. Persisters are phenotypic variants within the bulk population, but genetically identical (4, 17, 18). As a consequence, persisters can transform back into antibiotic-susceptible bacteria and, after the antibiotic pressure is removed, reconstitute a population that displays an antibiotic-tolerance identical to the starting culture (4, 18). Persister cells seem to be a universal feature of clonal life forms. Not only are they identified in many, if not all, bacterial species that were studied, also eukaryotic cancer cell lines and yeast populations contain drug-tolerant phenotypic variants (18–20). Many studies indicate the clinical relevance of persistence (7, 18, 21–23). Bartell *et al*. (2020) observed a link between high persister variants of *Pseudomonas aeruginosa*, long-term establishment of *P. aeruginosa* in the cystic fibrosis lung environment and treatment failure (24). Van den Bergh *et al* (2022) demonstrated the role of metabolic homeostasis, and more specifically of the respiratory complex I, as an important promotor of antibiotic persistence *in vitro* (25). Similarly, Zampieri *et al*. (2021) indicated the importance of the respiratory and fermentative metabolism in the preservation of heterogeneity within bacterial cultures and in adaptation of bacteria to environmental changes (26). Another clinical important consequence is that persistence is a driver towards the development of antibiotic resistance (18). Clearly, persisters constitute a viable pool that facilitate resistance development by prolonging the presence of viable bacteria during antibiotic treatment (27, 28), but various other mechanisms were suggested (6, 8, 10, 27, 29). For example, Windels *et al*. (2019) and Huo *et al*. (2022) identified that the increased mutation rate in high-persistent strains promotes evolution towards antibiotic resistance (6, 30) and Levin-Reisman *et al*. (2019) indicated the role of epistasis between antibiotic persistence and resistance mutations (29).

The presence of antibiotic-tolerant persisters in *S. pneumoniae* has not been investigated to date. In part, the lack of understanding persistence in *S. pneumoniae* stems from the self-limiting nature of this bacterium *in vitro* (31). Two suggested causes of the fast decrease in survival after entering the stationary phase are the enzymes pyruvate oxidase (SpxB) and autolysin (LytA) (32, 33). Pyruvate oxidase is the major producer of H_2_O_2_ as a by-product of the aerobic metabolism of *S. pneumoniae*, but *S. pneumoniae* lacks the neutralizing enzyme catalase which leads to *in vitro* death through an accumulation of H_2_O_2_ (32, 34–36). Autolysin, a cell-wall bound amidase that breaks down peptidoglycan, induces *in vitro* autolysis in stationary phase cultures (33, 37–39). Another reason why *S. pneumoniae* persisters might have been ignored can be the acute nature of *S. pneumoniae* infections (1). Antibiotic-tolerant persisters are mostly connected with recurrent and chronic infections and the role of persisters in acute infections is not clear (7, 18). Nonetheless, *S. pneumoniae* is also, albeit to a lesser extent, the causative agent of chronic diseases, like chronic endobronchial infections in children (40–42) and it can reside in biofilms in the middle ear of children causing recurrent and chronic otitis media (43–46). The role of persister cells, both in acute and chronic pneumococcal infections, needs to be elucidated in order to gain a better understanding in how *S. pneumoniae* evades elimination by antibiotic treatment (7, 18, 47).

In this study, we made a broad inquiry on the presence and behaviour of persister cells in the populations of diverse *S. pneumoniae* isolates. We succeeded in obtaining stable long-living *in vitro* cultures using specific growth conditions which allowed us to set-up prolonged antibiotic-induced killing studies without confounding the results with the self-limiting nature of *S. pneumoniae*. Using these killing studies together with heritability assays, the golden standard assays to determine persistence (4, 18), we proved the presence of high numbers of persister cells in reference strain D39 cultures. Lastly, we detected the presence of persister cells in a variety of *S. pneumoniae* strains, including clinical isolates, demonstrating that persistence is widely present and highly variable in *S. pneumoniae*. Our study is the offset for persistence studies in *S. pneumoniae* and will lead to better insights in the role of persistence in acute, chronic and recurrent *S. pneumoniae* infections. In turn, a better understanding of the escape mechanisms of *S. pneumoniae* will finally lead to improved therapeutic options.

## RESULTS

### Specific growth conditions allow long-living *in vitro* cultures of *S. pneumoniae*

To study persistence, prolonged *in vitro* antibiotic-induced killing studies are required. Especially when examining antibiotic survival in stationary phase, long-living cultures are needed and any confounding effects of mortality through the self-limiting nature of *S. pneumoniae* must be avoided. In order to prevent *in vitro* death in absence of antibiotics and to obtain a stable bacterial culture, we followed two routes targeting the suggested effectors of self-limitation in *S. pneumoniae*. First, we added catalase to neutralize the produced H_2_O_2_, we constructed a *spxb* knockout mutant to inhibit the expression of pyruvate oxidase or we applied hypoxic incubation (5% CO_2_ – 0.1% O_2_ – 94.9% N2) to inhibit the pneumococcal aerobic metabolism and thus the production of H2O_2_ by pyruvate oxidase (32, 34). Second, we used choline chloride supplementation to prevent the binding of autolysin to the cell wall or we used a *lyta* knockout mutant to inhibit the expression of autolysin (33, 37–39).

Despite the common knowledge and the regular growth in exponential phase (31, 48), a phase of significant killing after 8-48 hours of incubation was unavoidable in the commonly used growth media Brain Heart Infusion broth (BHI) and Todd-Hewitt broth supplemented with 0.5% Yeast extract (THY), with or without applying the different strategies to counteract pyruvate oxidase or autolysin *(supplementary data figure 1)*. While a section of the stationary phase was reasonably stable (5-20 fold reduction in bacterial concentration from 16 to 24h) when adding catalase or using a *lyta* knockout mutant, the bacterial concentration was nevertheless reduced up to 1,000 fold before reaching such stable period. Surprisingly, when using a less common growth medium, Mueller-Hinton broth supplemented with 5% Lysed horse blood (MHL), *in vitro* self-limitation is mostly absent during 32 hours of incubation (Figure 1). Only after 32 hours, a strong death phase occurs with a 10,000 to 1,000,000 fold reduction in bacterial viability. Also here, counteracting pyruvate oxidase and autolysin by applying the proposed strategies did not significantly impact survival (Figure 1). Therefore, using MHL seems to be the optimal liquid growth medium to obtain a stable long-term bacterial culture.

**Figure 1:**
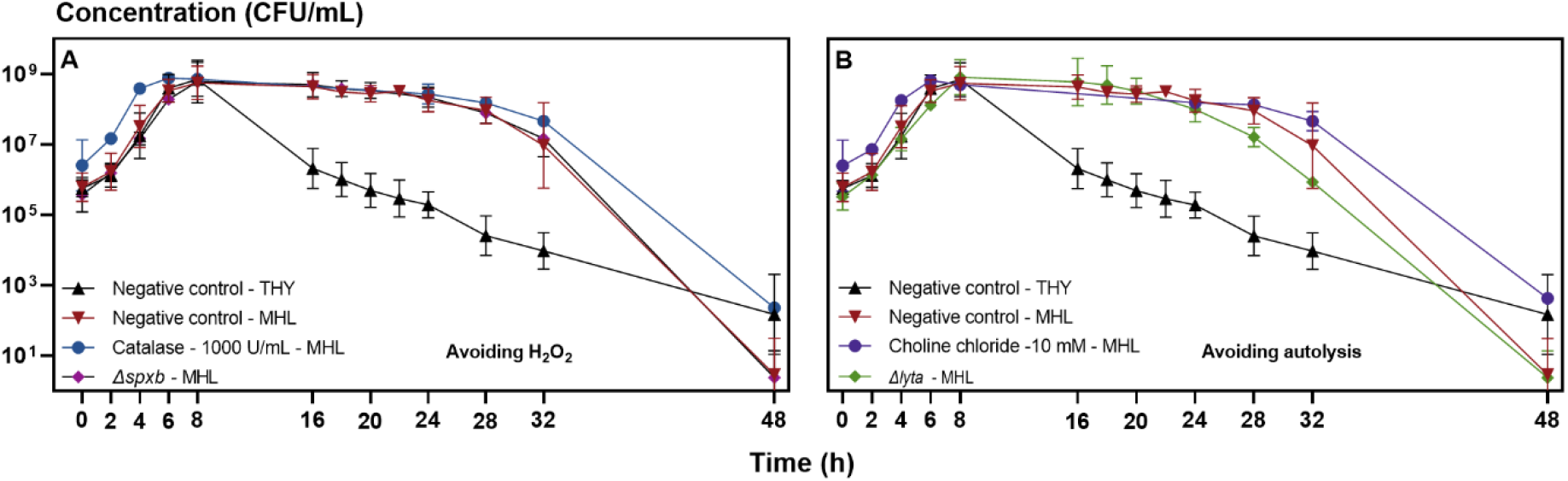
MHL abolishes the self-limiting *in vitro* nature of *S. pneumoniae* regardless of the strategies adopted to avoid H_2_O_2_ or autolysis. We compared the effect of different strategies to counteract the effects of pyruvate oxidase (A) or autolysin (B) in planktonic growth curves of *S. pneumoniae* D39 in MHL (Mueller-Hinton broth supplemented with 5% Lysed horse blood). The planktonic growth curve of *S. pneumoniae* in THY (Todd-Hewitt broth supplemented with 0.5% Yeast extract) is repeated for visual comparison. The different strategies didn’t prolong the survival over 48 hours of *S. pneumoniae* compared to MHL alone, but survival is higher for all conditions compared to THY. The experiments were performed in duplicates or triplicates and each value is presented as the mean ± standard deviation (n ≥ 2).

Culture conditions are important for *S. pneumoniae* survival *in vitro* and the use of the liquid growth medium MHL results in stable survival. Therefore, we optimized a model based on MHL as growth medium (Figure 2). To validate the model for survival over 24 hours, growth curves were obtained for five *S. pneumoniae* strains (Figure 3). Strains were either commonly used lab strains (TIGR4, ATCC49619 and R6) or lab strains from previous studies (85 and 88) (49). While these strains show small differences in lag phase, growth rate and maximal bacterial concentration, overall survival over 24 hours is stable among all strains (Figure 3). In conclusion, the optimized long-living model results in a stable bacterial culture until 24 hours of growth which enabled us to execute prolonged time-killing experiments without the self-limiting nature of *S. pneumoniae* as confounding factor. We applied this model in all further experiments described in this manuscript.

**Figure 2:**
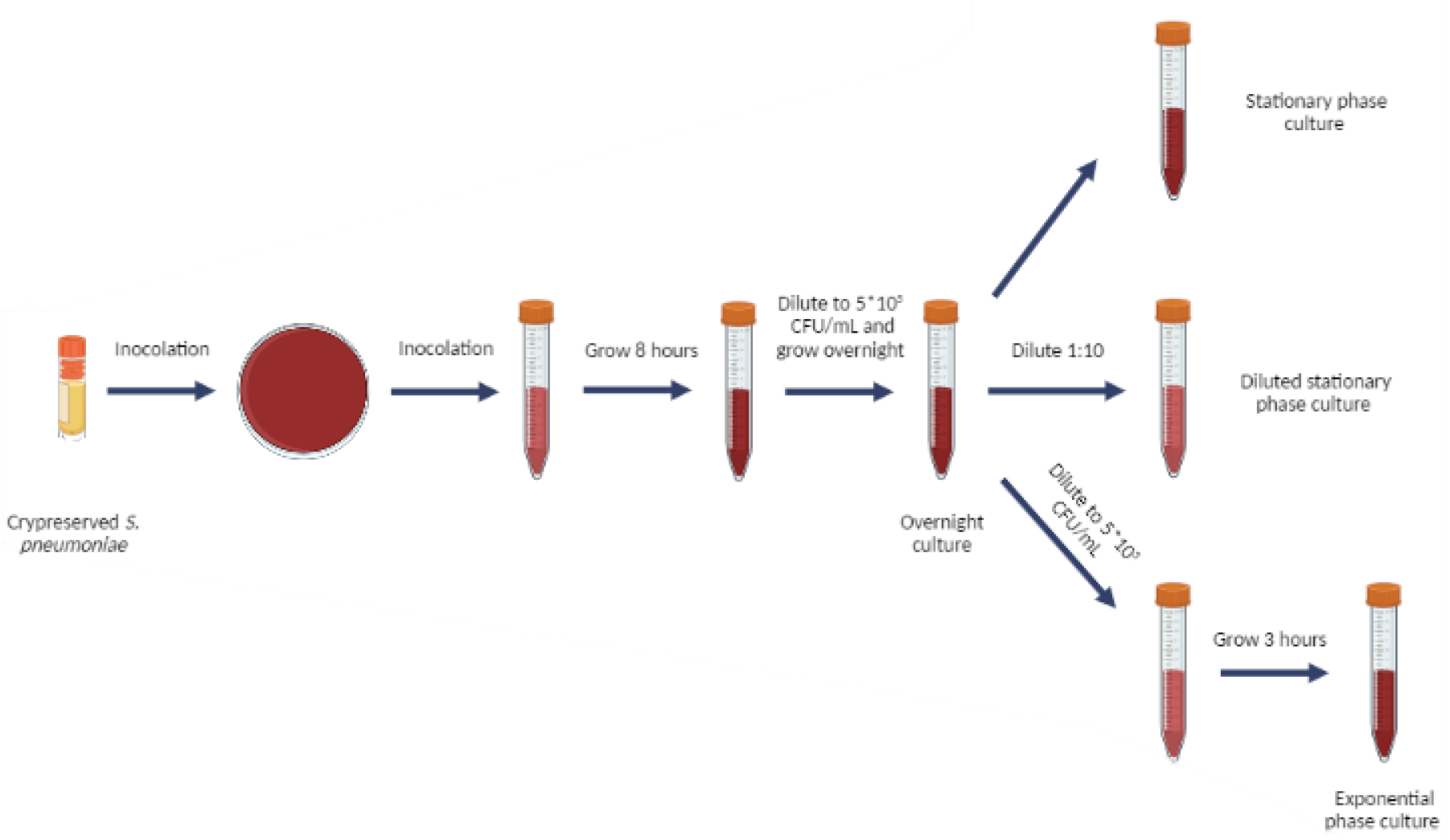
A schematic overview of the long-living *in vitro* culturing protocol that results in stable bacterial cultures. Cryopreserved *S. pneumoniae* are plated on a blood agar plate followed by inoculation in a tube with MHL. After 8 hours of static incubation, the culture is diluted to 5*10^5^ CFU/mL and grown overnight. The overnight culture is either directly used as a stationary phase culture, diluted 1:10 in fresh MHL to act as a diluted stationary phase culture or is diluted to 5*10^5^ CFU/mL in fresh MHL and grown 3 hours to obtain an exponential phase culture.

**Figure 3:**
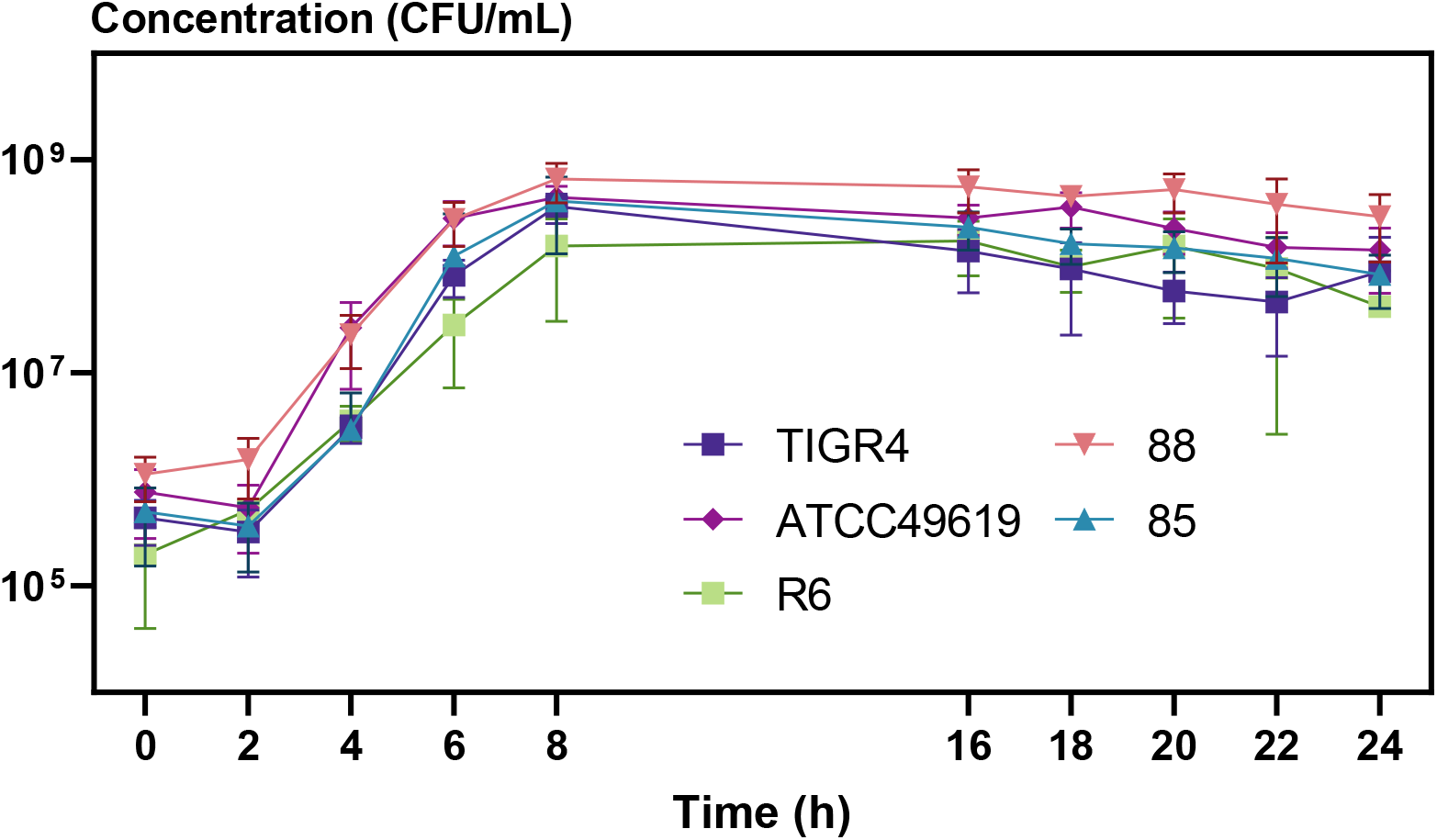
Various *S. pneumoniae* lab strains show robust growth dynamics for up to 24 hours of incubation using the optimized long-living *in vitro* model. Planktonic growth curves in function of time of *S. pneumoniae* TIGR4, ATCC49619, R6, 85 and 88 show small differences in lag phase, growth rate and maximal bacterial concentration, but a stable survival over 24 hours. The experiments were performed in triplicates and each value is presented as the mean ± standard deviation (n = 3).

### Persisters are widely present in *S. pneumoniae* reference strain D39 cultures and highly dependent on growth phase and type of antibiotic

To study persister cells, the concentration of the applied bactericidal antibiotics needs to be well above the minimal inhibitory concentration (MIC) to invoke killing of sensitive cells. Reference strain D39 is sensitive to amoxicillin, cefuroxime, moxifloxacin and vancomycin - clinically relevant antibiotics of various classes - according to the EUCAST breaking points (*supplementary data table 1*). To evaluate antibiotic-induced killing, applying an excess of such MIC concentrations is thus straightforward in further experiments. Along with the selection of antibiotics and their concentration, we tested different growth phases for treatment of *S. pneumoniae* with the antibiotics. Three growth conditions are frequently used to score persister levels, the stationary phase, the diluted stationary phase and the exponential phase (50–52). The protocol we used to obtain these different growth phases is described in Figure 2. Briefly, cryopreserved *S. pneumoniae* were plated on a blood agar plate followed by inoculation in a tube with MHL. After 8 hours of static incubation, the culture was diluted to 5*10^5^ CFU/mL and grown overnight. The overnight culture was either directly used as a stationary phase culture, diluted 1:10 in fresh MHL to act as a diluted stationary phase sample or was diluted to 5*10^5^ CFU/mL in fresh MHL and grown 3 hours to obtain an exponential phase culture.

**Table 1:**
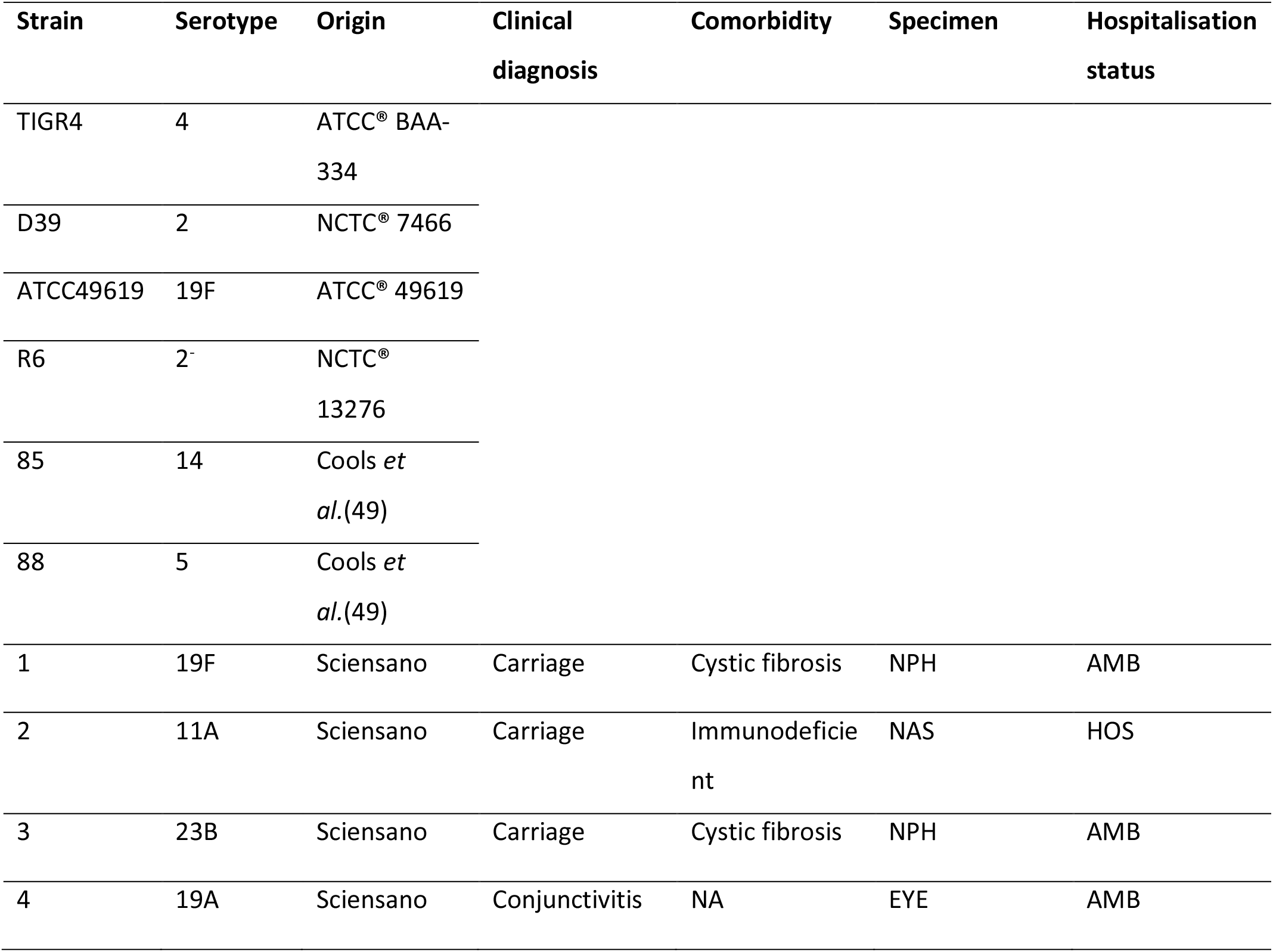
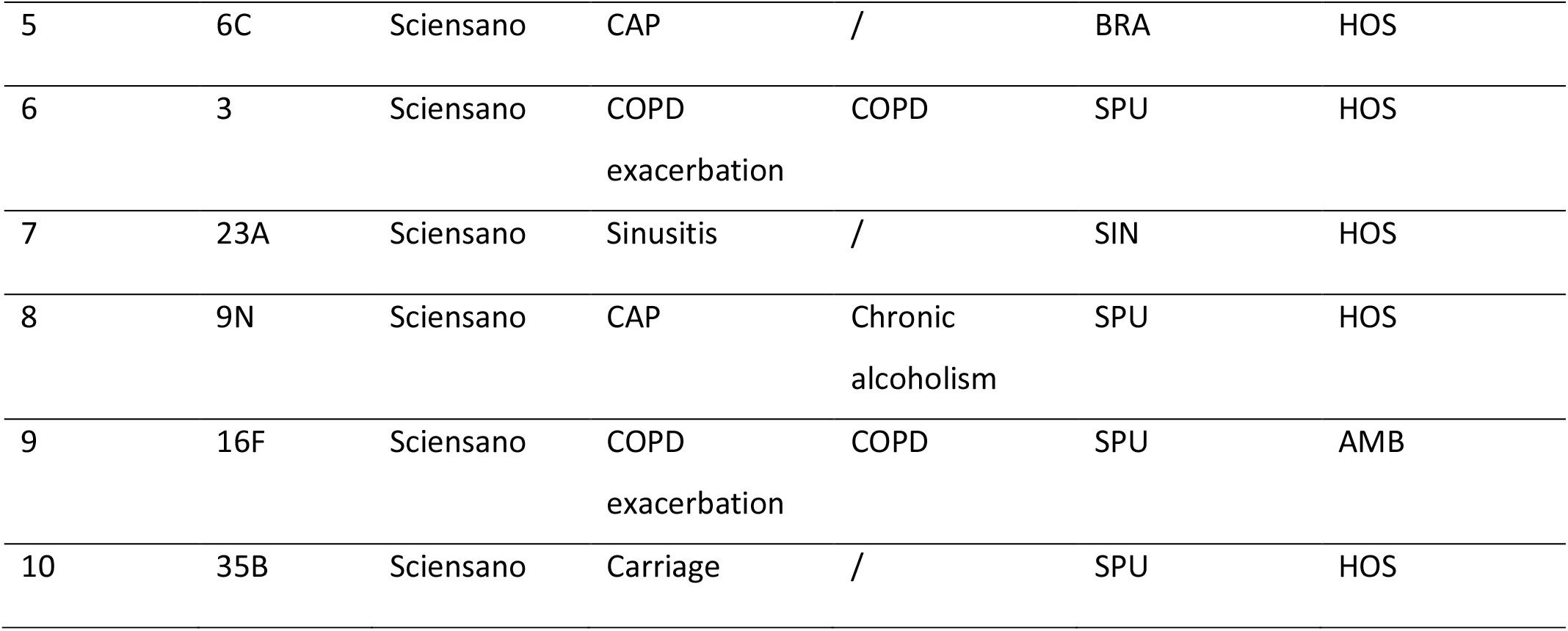
*S. pneumoniae* strains used during this study. Abbreviations: NA = not available, CAP = community acquired pneumoniae, COPD = chronic obstructive pulmonary disease, NPH = Nasopharyngeal aspirate/swab, NAS = nasal swab, BRA = endotracheal/bronchial aspiration, SPU = sputum, SIN = sinus, AMB = ambulatory, HOS = hospitalized.

To evaluate the minimal dose needed to kill sensitive *S. pneumoniae* D39 cells within 5 hours, we obtained dose-dependent kill curves by treating *S. pneumoniae* with increasing antibiotic concentrations, i.e. 20-, 100- and 200-fold the MIC. Stationary phase cultures proved insensitive to any of the used antibiotics, even at the highest dose, but treatment of diluted stationary phase samples resulted in significant killing of sensitive cells at or above a concentration of 20-fold the MIC (Figure 4). The independence on antibiotic concentration, once a sufficient dose is reached, is a typical observation indicating a role for persistence while strong correlations would point towards antibiotic resistance as underlying cause of survival (4). For the remainder of our work, we applied concentrations of 100-fold the MIC.

**Figure 4:**
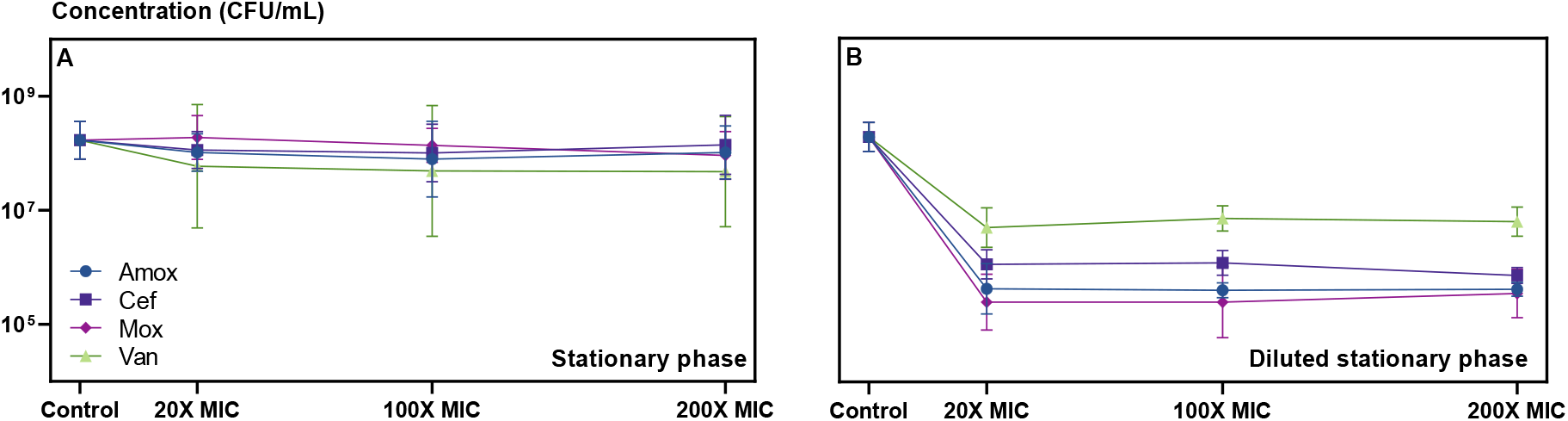
*S. pneumoniae* D39 in stationary phase is insensitive to antibiotic treatment while a dose of 20-fold the MIC is sufficient to kill sensitive cells in a diluted stationary phase culture. Dose-dependent kill curves with amoxicillin (amox, blue dots), cefuroxime (cef, dark purple squares), moxifloxacin (mox, light purple diamonds) and vancomycin (van, green triangles) of planktonic *S. pneumoniae* D39 in (A) stationary phase or (B) diluted stationary phase samples. Antibiotic treatments lasted for five hours before enumerating survivors. Applied concentrations were 20-, 100- and 200-fold the MIC (respectively 0,14; 0,70 and 1,40 µg/mL for amoxicillin, 0,54; 2,7 and 5,4 µg/mL for cefuroxime, 5,8; 29 and 58 µg/mL for moxifloxacin and 8,8; 44 and 88 µg/mL for vancomycin). The experiments were performed in triplicates and each value is presented as the mean ± standard deviation (n = 3).

### *Streptococcus pneumoniae* D39 cultures contain high numbers of persisters

To investigate the presence of persister cells, we followed the survival of cells in function of time during antibiotic treatment. These so-called time-kill curves should show a single rate of killing (uniphasic pattern) if the bacterial culture is fully susceptible to the antibiotic and lacks any subpopulation with increased tolerance (persister cells). However, if a subpopulation of antibiotic-tolerant persister cells is present within the susceptible population, we expect distinctly different killing rates to be apparent in the time-kill curves (biphasic pattern).

As stationary phase cultures did not show any killing (Figure 4), we performed these time-kill assays on diluted stationary phase and exponentially growing samples. Upon dilution of stationary phase cultures, antibiotic treatment kills 90-99.99% of the cells of strain D39 over an 8-hours period, depending on the used antibiotic (Figure 5). We observed similar killing after an 8-hours treatment of exponentially growing *S. pneumoniae* D39, but when treatment is prolonged to 24 hours, antibiotic treatment kills an additional 3 orders of magnitude of the exponentially growing cells (Figure 5). Mathematical analyses of the entire dataset, with a global model containing a condition-dependent structure, showed that the biphasic killing model is superior in describing the data compared to the uniphasic model (ANOVA (F-test), p = 1,58e-84; *supplementary data table 2*), which implies that the sensitive *S. pneumoniae* D39 population contains persister cells. When each condition (growth phase x antibiotic) is analyzed separately, the biphasic model is preferred over the uniphasic model for describing the data from all conditions (p ≤ 0.05), except for data from treatment with cefuroxime (p = 0.1641) and vancomycin (p= 0.1074) in the diluted stationary growth phase, indicating that including a second killing rate does not improve the models although various test statistics (AIC, BIC and LogLik) are approaching significance (*supplementary data table 2*). Overall, we detected relatively high persister levels, ranging from 13.74 to 60.08%, for the diluted stationary growth phase compared to the lower levels, ranging from 0.02 to 0.5%, in the exponential growth phase. The killing rates of persisters cells (0.25-0.58 h^-1^) are comparable between the different conditions and are 3- to 8-fold lower than the killing rates of normal cells (0.89 – 3.78 h^-1^) (*supplementary data table 3*).

**Figure 5:**
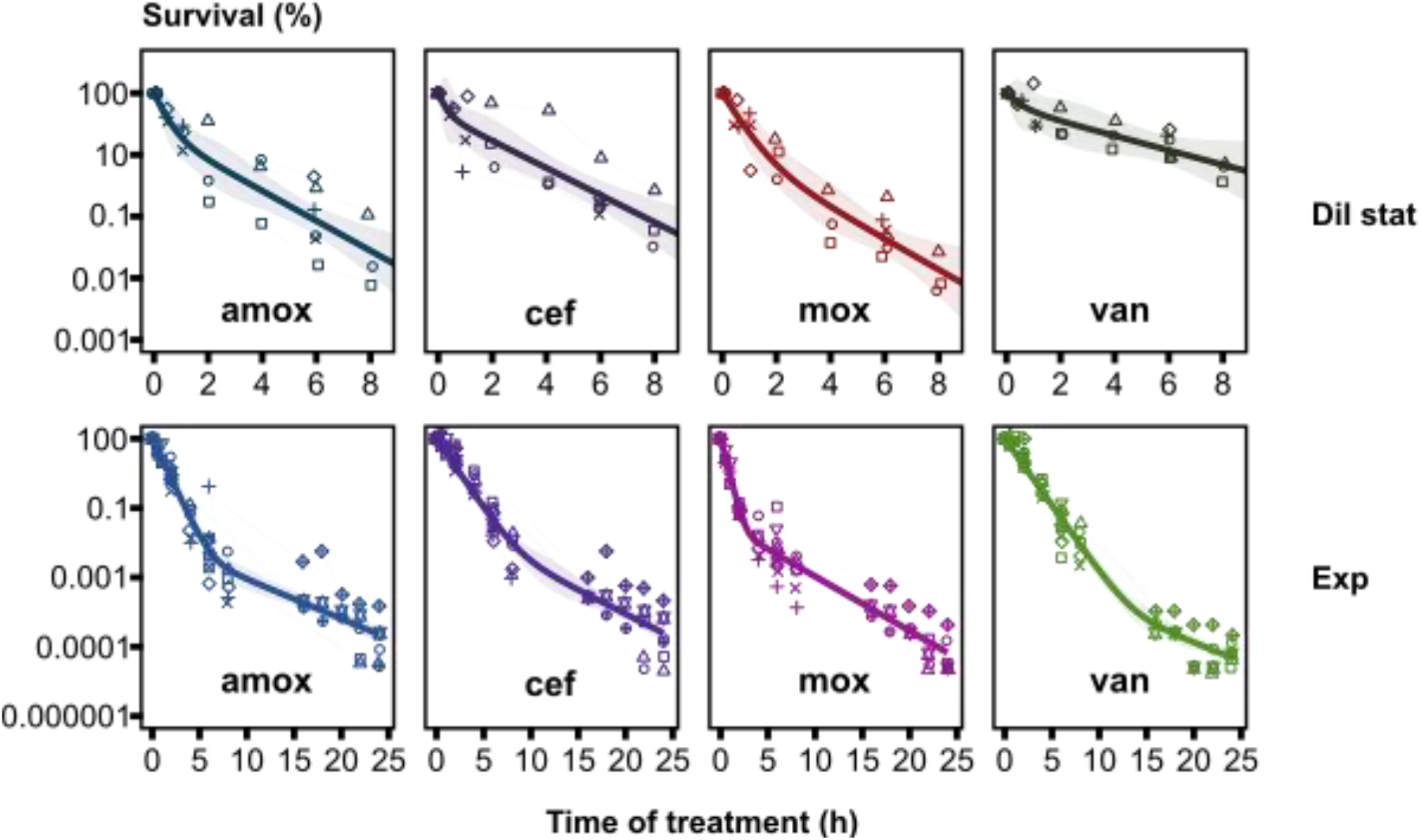
Biphasic killing pattern upon antibiotic treatment indicates presence of persister subpopulations in *S. pneumoniae* D39 cultures. Fitting of a non-linear fixed-effect model to log-transformed survival data upon treatment with amoxicillin (amox), cefuroxime (cef), moxifloxacin (mox) and vancomycin (van) against *S. pneumoniae* D39. 1:10 diluted stationary phase (Dil stat) and exponential phase (Exp) bacteria were treated during 8 or 24 hours with the antibiotic (100-fold the MIC, 0.70 µg/mL for amoxicillin, 2.7 µg/mL for cefuroxime, 29 µg/mL for moxifloxacin and 44 µg/mL for vancomycin). Symbols show the individual repeats (time-point connected and in the same shape if coming from the same repeat) and bold lines show the fitted biphasic killing curves ± 95% CI (shades) (n ≥ 3).

### The antibiotic tolerant *S. pneumoniae* persisters are transient and non-heritable

While biphasic killing patterns are the golden standard to identify persistence, theoretically, such surviving cells could still be the result of emerging resistance or of mutants that display an increased population-wide tolerance. To confirm the presence of persisters, we performed so-called heritability assays (4). Here, we re-tested some of the surviving clones of the initial time-kill assay in a subsequent round of treatment. If *S. pneumoniae* would be resistant, the MIC value would be increased and if the persister phenotype would be inherited and passed to the entire population of daughter cells, an increased survival would be observed during the subsequent killing-assays. During these subsequent killing-assays, we observed a similar survival of randomly selected clones that survived the initial killing assay (i.e. supposed persisters) (Figure 6), a similar killing dynamic pattern (*supplementary data figure 2)* and MIC values that remained unchanged (*supplementary data table 4*) compared to the original culture. Thus, the surviving cells that we observed are genuine persister cells showing only a transient antibiotic tolerance as regrown cultures show similar characteristics to the culture of origin.

**Figure 6:**
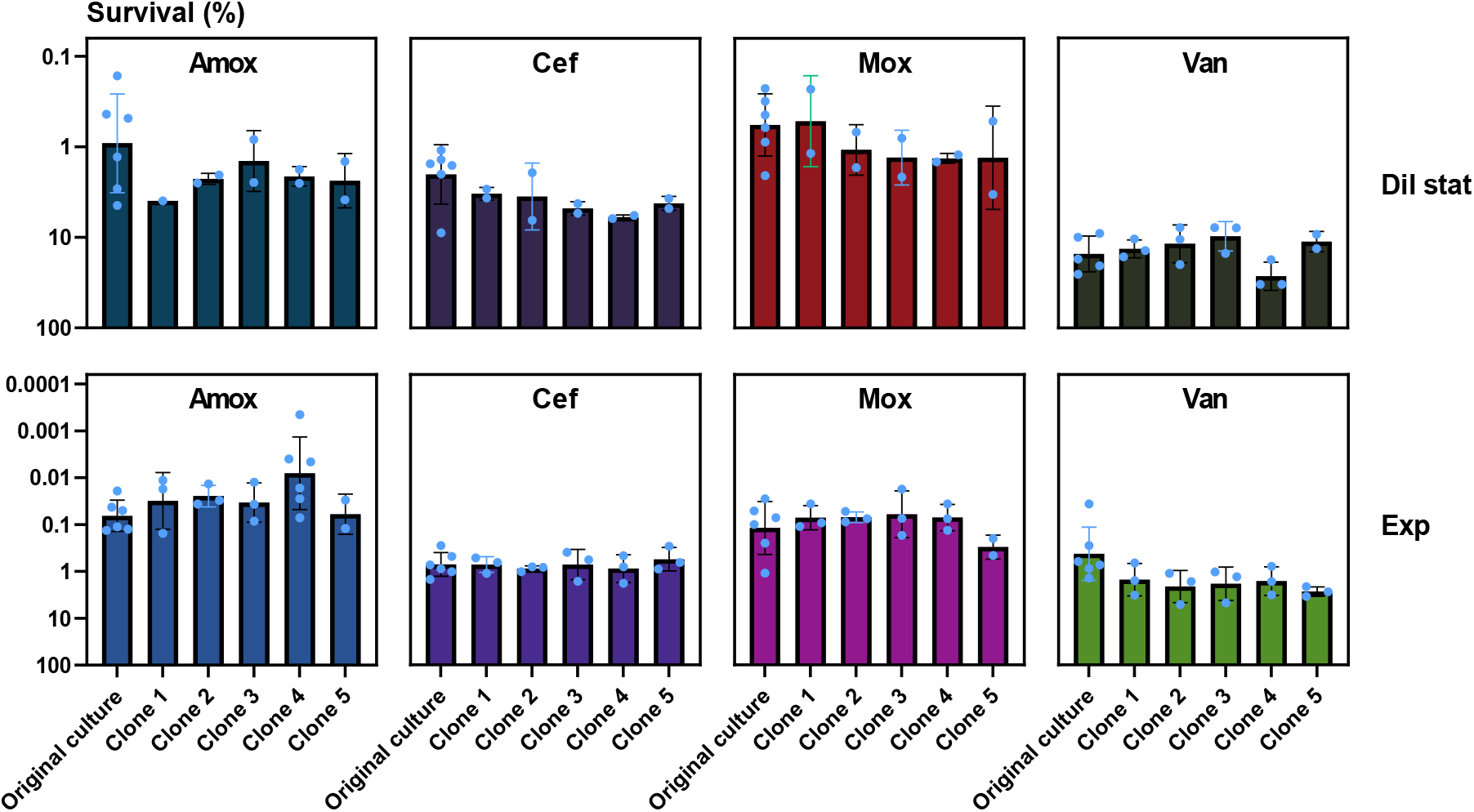
The antibiotic tolerance of surviving *S. pneumoniae* cells is transient and non-deterministically inherited by daughter cells. AB-tolerant *S. pneumoniae* D39 clones were recovered after 6 (Dil stat) or 18 (Exp) hours of treatment during the initial time-kill assay, regrown without antibiotic and preserved at -80°C. On these clones arising from potential persister cells, survival was determined after 6 hours of antibiotic treatment with amoxicillin (amox), cefuroxime (cef), moxifloxacin (mox) and vancomycin (van) in the diluted stationary (Dil stat) or the exponential growth phase (Exp). Survival of the randomly selected clones was similar to the original culture (one-way ANOVA). The experiments were performed in triplicates and each value is presented as the mean ± standard deviation (n = 3).

### Persistence is both prevalent as well as highly variable among *S. pneumoniae* clinical isolates

Having established the presence of persisters in strain D39, we wondered whether persistence is a wide-spread phenotype among *S. pneumoniae* strains and how this phenotype varies within the *S. pneumoniae* species. To answer this questions, we obtained a set of 10 clinical isolates (CI 1-10) representing 10 different serotypes with known origin in addition to the already available strains (D39, TIGR4, ATCC49619, R6, 85 (49) and 88 (49)). Clinical isolates were either isolated from patients suffering pneumococcal disease (CI 4-9) or from patients that were only carriers (CI 1-3 and CI 10). The infections that were caused by these strains, were conjunctivitis (CI 4), community acquired pneumonia (CAP) (CI 5 and CI 8), sinusitis (CI 7) or the CI was isolated from patients during a COPD exacerbation (CI 6 and CI 9). 3 out of 4 patients that carried *S. pneumoniae* suffered from comorbidities like cystic fibrosis or immune deficiency and 3 out of 6 patients that suffered from pneumococcal disease had comorbidities like COPD or chronic alcoholism (Table 1).

All strains were susceptible to amoxicillin, cefuroxime, moxifloxacin and vancomycin, except for strain 85 which displayed cefuroxime resistance and CI 7 which displayed minor moxifloxacin resistance according to the EUCAST breaking points (*supplementary data table 5*). To screen for persister formation, *S. pneumoniae* strains were challenged with either amoxicillin or vancomycin (100-fold the MIC) during 8 hours in the diluted stationary or exponential growth phase in order to determine survival (Figure 7). Overall, we observed higher survival fractions for *S. pneumoniae* strains treated in the diluted stationary phase compared to treatment in the exponential growth phase. Survival after treatment with vancomycin was higher than after challenging *S. pneumoniae* with amoxicillin in the diluted stationary phase, but survival fractions were comparable after treatment of exponentially growing bacteria. When we compared the survival of *S. pneumoniae* within a condition (growth phase x antibiotic), we detected large variations between strains ranging over four orders of magnitude for each condition. Strain R6, an unencapsulated reference strain, displayed the overall highest survival while CI 10, an isolate from carriage, displayed the lowest. For all conditions, survival was significant higher for CI 1 compared to CI 10, which were both carriage isolates (one-way ANOVA, p ≤ 0.01). No significant differences were detected, in any condition, between the isolates from acute infections (p > 0.05), except for CI 4 which had a significant lower survival than CI 7 after treatment with vancomycin in the diluted stationary phase (p = 0.0461). Survival, and thus tolerance levels, are highly variably in *S. pneumoniae* strains from different sources.

**Figure 7:**
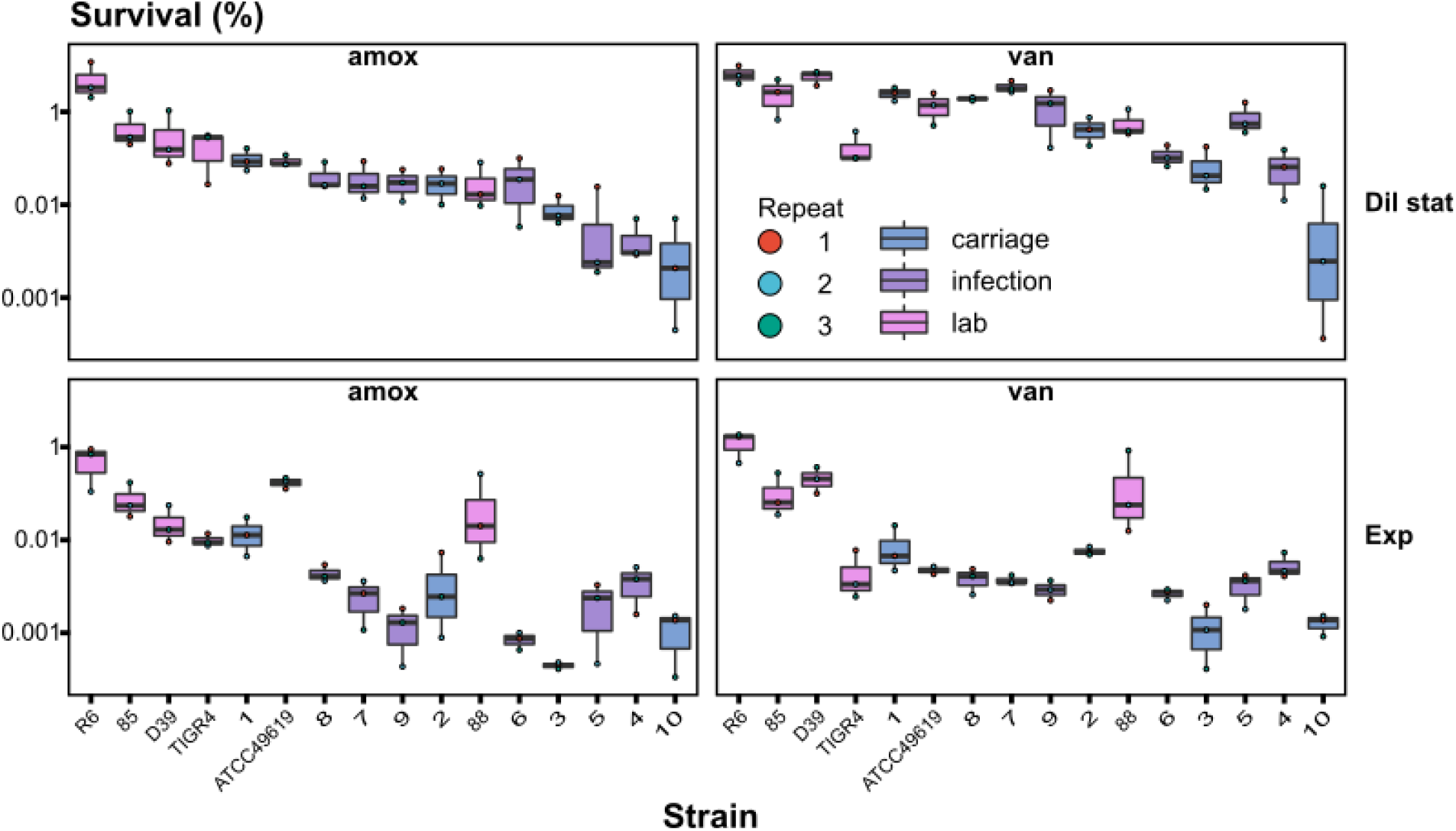
Persistence is widely present and highly variable in *S. pneumoniae* strains from different sources. Survival is highly variable in *S. pneumoniae* strains after challenge with 100-fold the MIC of amoxicillin (amox) or of vancomycin (van). An 8-hours antibiotic treatment was started after a 1:10 dilution of stationary phase bacteria (Dil stat) or after 3 hours of growth (exponentially growing bacteria; Exp). Strains are ordered based on their survival after treatment with amoxicillin in the diluted stationary phase. Log-transformed data are shown as a boxplot of the mean ± standard deviation for each strain. The experiments were performed in triplicates and each individual measurement is given as a dot according to the repeat (n = 3).

Given the strong variation between strains, we wondered whether some strains show persistence specific in one condition or whether survival levels of these strains can be correlated between different conditions. We observed strong correlations between the growth phases (diluted stationary and exponential) and the antibiotics (amoxicillin and vancomycin) (Figure 8 and *supplementary data figure 3*). Surprisingly, we also detected a small positive correlation (Pearson correlation, R² ranging from 0.032 – 0.26) between increased growth and increased survival, which contradicts the common believe that slow growth induces persister formation (*supplementary data figure 3*). After screening different *S. pneumoniae* strains for persister formation, we can conclude that persisters are widely present in *S. pneumoniae* cultures from different sources (lab strains, clinical isolates from infection and carriage) with strong variations between strains within a condition.

**Figure 8:**
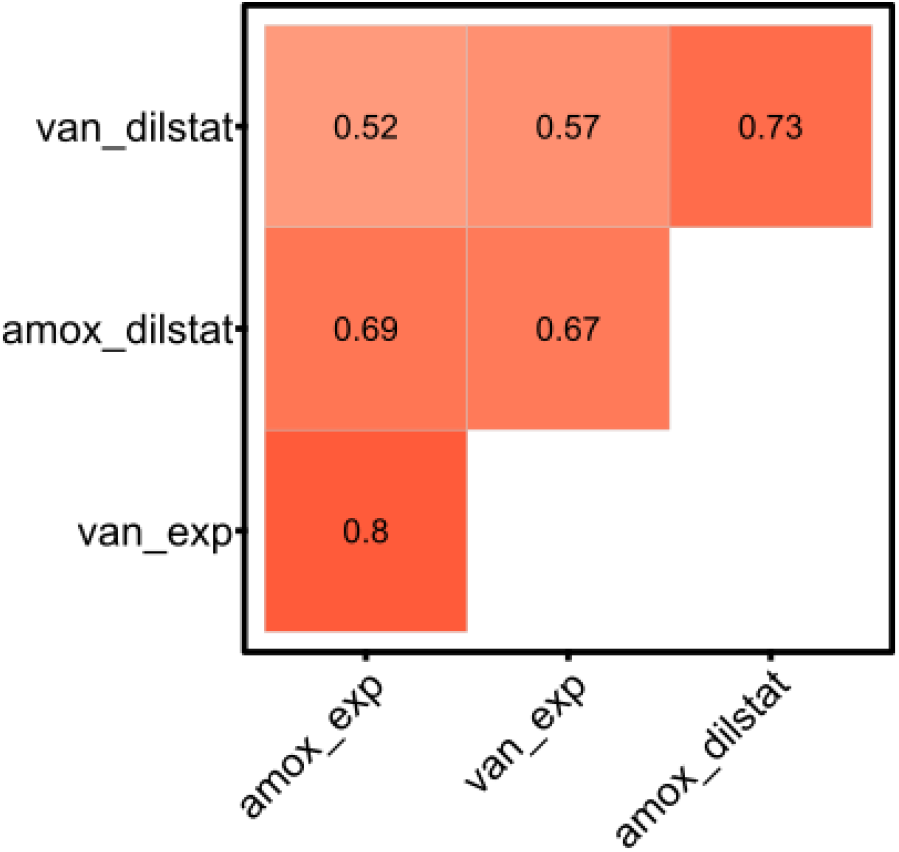
Correlation analysis of survival between different conditions show strong correlations between antibiotics (amoxicillin and vancomycin) and growth phases (diluted stationary and exponential growth phase). Correlation matrix between the survival fractions in 4 different conditions: treatment with amoxicillin (amox) or vancomycin (van) in the diluted stationary (dilstat) or exponential (exp) growth phase. Pearson correlation coefficients (R) are given for each correlation.

## DISCUSSION

Our study presents a broad characterization of persistence in *S. pneumoniae*. We confirmed the hypothesis that persister cells are present in *S. pneumoniae* cultures by finding strong indications of a biphasic killing pattern, the hallmark of persistence, after treatment with four clinically relevant antibiotics with different modes of action. Additionally, the surviving persisters in *S. pneumoniae* show no inheritable resistance, as tolerance to the antibiotics was not passed from the initial persister cell to the subsequent generation during the heritability assays and the MIC values remained unchanged. Finally, a set of clinical isolates was screened for persistence. Here, we identified large variations in persistence among different strains, suggesting persistence is a general trait in *S. pneumoniae* cultures and the phenotype is controlled by a broad range of genetic elements.

Most insights in persistence have been gathered using Gram-negative bacteria, like *E. coli, S. typhimurium* and *P. aeruginosa* (8, 10, 11, 18). However, different studies already indicated a role for persistence in Gram-positive bacteria as well (18, 53–55), and more specifically in various reports on streptococcal species (18, 56). Persisters in *S. mutans*, a cariogenic oral bacterium, are tolerant to a wide variety of antibiotics (57), but also to a dental caries defensive agent, dimethylaminododecyl methacrylate (DMADDM) (58), and to other antibacterial monomers used in dental medicine (59). As in non-streptococcal bacterial species, toxin-antitoxin systems are involved in the formation of persister cells in *S. mutans*, as well as the quorum-sensing peptide CSP (competence stimulating peptide) (57, 60). Similarly, antibiotic-tolerant persisters were identified in the zoonotic pathogen *S. suis* by Willenborg *et al*. (2014) (61) and in the opportunistic human pathogen *S. faecalis* as early as 1979 by *Soriano et al*. (1979) (62). Additionally, persistence in *S. pyogenes* was observed by Wood *et al*. (2005) in stationary phase *in vitro* cultures (63) and Martini *et al*. (2021) detected persister cells in *S. pyogenes* biofilms treated with antimicrobials (64). Despite the various reports on persistence in other species of the *Streptococcus* genus, very little is known about antibiotic tolerance, and more specific about persistence, in the Gram-positive bacterium *S. pneumoniae* (13–16).

*S. pneumoniae* is well-known to cause acute infections while antibiotic tolerance and persistence are mostly connected with recurrent and chronic infections (2, 7). Nonetheless, we expected to find persister cells as persistence is identified in many, if not all, bacterial species that were studied and *S. pneumoniae* causes, to a lesser extent, also recurrent and chronic infections (41, 42, 45, 65). After mathematical analyses of the killing data of *S. pneumoniae* D39, we detected a biphasic killing pattern, which indicates the presence of persister cells. A major advantage of our approach is that we determined persister fractions, and killing rates, by mathematical analysis based on kill curves over a prolonged treatment period which enabled us to take into account the killing pattern rather than determine the persister fraction based on a single timepoint. The characteristics of persistence differed between growth phases and antibiotics. Persister levels were vastly higher in diluted stationary phase cultures than for exponentially growing bacteria for all examined strains, as we expected, because persistence is mostly linked to dormancy and bacteria become less metabolically active when they enter the stationary phase (21, 43, 66). Interestingly, treatment of diluted stationary phase cultures with moxifloxacin, a fluoroquinolone that targets the DNA synthesis of bacteria and is less dependent on cell growth than β-lactams, resulted in the lowest level of persisters (13,74%) in diluted stationary phase cultures (47, 67, 68). Overall, we observed higher persister fractions than previously described, but a potential explanation is the difference in approach, we used mathematical analysis based on prolonged time-kill curves rather than on survival at a single timepoint. If we would determine the persister fraction based on survival fractions after a fixed period of treatment, for example after 8 hours, fractions approximate 0.01-1% as is described for *S. mutans* or *E. coli* stationary phase cultures (51, 57). Altogether, we detected that persisters are widely present in *S. pneumoniae* reference strain D39 cultures and that *S. pneumoniae* persistence is highly dependent on growth phase and the type of antibiotic.

Finally, a set of *S. pneumoniae* strains was screened for persister formation. In total, 16 strains were screened, including 6 lab strains (D39, TIGR4, ATCC49619, R6, 85 (49) and 88 (49)) and 10 clinical isolates (CI 1-10). This set of *S. pneumoniae* strains is diverse with different serotypes and from different origins (lab, infection or carriage). Survivors were detected for all strains in various levels, ranging from 0.001 to 10% surviving cells, which showed that persistence is widely present, but also highly variable in *S. pneumoniae* cultures. Stewart *et al*. (2012) and Hofsteenge *et al*. (2013) detected comparable variation in survival after antibiotic treatment of a set of natural and environmental *E. coli* isolates, respectively (69, 70). Similarly, Barth *et al*. (2013) observed a high heterogeneity of persister cell formation among *Acinetobacter baumannii* isolates (71). We analyzed the survival data for correlations (Pearson) between the growth phases and antibiotics. The strong correlation between the growth phases indicate that these conditions are affecting persistence in a similar way, potentially because they both induce growth and metabolic activity. Furthermore, survival data for both types of antibiotics, amoxicillin and vancomycin, correlate well, potentially as a consequence that they both target the bacterial cell wall synthesis. Surprisingly, we saw a slight positive correlation between how fast the strains grow and how well they survive antibiotic treatment, especially for exponentially growing bacteria. These correlations imply that if the bacteria grew faster, they survived antibiotic treatment better. This was unexpected, as persistence is linked to dormancy of bacterial cells and we expected that if cells were less actively dividing and less metabolically active, they would survive antibiotic treatment better. However, different studies state that global metabolic dormancy is not solely responsible for tolerance (72–76). For example, Stapels *et al*. (2018) and Peyrusson *et al*. (2020) demonstrated the presence of non-dividing, but metabolically active *Salmonella* and S*taphylococcus aureus* persisters, respectively, during intracellular infections (77, 78) and Goneau *et al*. (2014) stated that antibiotic tolerance is caused more likely by selective target inactivation than by global metabolic dormancy in uropathogens (79). The screening of a variety of *S. pneumoniae* strains showed that persistence is widespread and diverse among *S. pneumoniae* cultures suggesting that a broad range of genetic elements are controlling the phenotype. Further screening of clinical isolates is necessary to be able to correlate persistence with serotype and origin, but also to study the role of *S. pneumoniae* persisters in treatment outcome.

With this study, we made a broad characterization of persistence in *S. pneumoniae*. First, we obtained long-living *in vitro* cultures of *S. pneumoniae* eliminating its self-limiting nature. Secondly, we detected the presence of a biphasic killing pattern after analyzing antibiotic-induced time-kill assays, the hallmark of persistence, and we proved that *S. pneumoniae* persistence is transient and not-heritable. Finally, we observed persister cells in *S. pneumoniae* strains from different origins which showed that persistence is widely distributed and highly variable among *S. pneumoniae* isolates. Together, our work advocates for higher interest for persistence in *S. pneumoniae* as a contributing factor for therapy failure and resistance development. Future studies should gain better insights in the mechanisms of persister formation and improve knowledge about the clinical relevance of pneumococcal persisters. Therefore, further screening of clinical isolates and *in vivo* studies are required. The ultimate goal is to gain better insights in the role of persistence in acute, recurrent and chronic *S. pneumoniae* infections what will hopefully lead to improved therapeutic options.

## MATERIALS AND METHODS

### Bacterial strains and growth conditions

*S. pneumoniae* strains used are listed in Table 1. *S. pneumoniae* was cultured statically in Brain-Heart Infusion broth (BHI) (Neogen), Todd-Hewitt broth (BD Biosciences) supplemented with 0.5% yeast extract (Gibco, THY), cation-adjusted Mueller-Hinton broth (Fluka) supplemented with 5% lysed horse blood (Oxoid, MHL) or on blood agar (BA) plates (Tryptic Soy Agar (Neogen) supplemented with 5% defibrinated sheep blood (Oxoid)) at 37°C in 5% CO_2_. Catalase (30 000 U/mL, MP biomedicals) or choline chloride (Sigma-Aldrich) was added when specified. Bacteria were grown in hypoxic conditions (5% CO_2_ – 0.1% O_2_ – 94.9% N2) in a hypoxystation (Whitley H35 Hypoxystation) when specified. *Escherichia coli* strain DH5-α was cultured shaking in Luria-Bertani broth (Lennox) (LB, Sigma-Aldrich) at 37°C, 175 rpm.

### Planktonic growth and enumeration of bacteria

Bacteria were grown in different media with or without catalase or choline chloride supplementation. At different time-points, samples were taken and the bacterial concentration was determined according to the viable plate count (VPC) method. Briefly, a 1:10 serial dilution (10^0^-10^−6^) was made in phosphate-buffered saline (PBS) in a 96-well plate. Three drops of 10 µL of a selection of dilutions was plated on BA and incubated for minimum 24 hours before colonies were counted and suspensions were enumerated.

### Long-living *in vitro* culturing

Bacteria from cryopreservation were plated on a blood agar plate and incubated for 24 to 72 hours followed by subculturing in a tube with MHL during 8 hours with a final concentration of 1*10^8^ CFU/mL. Then, bacteria were diluted to 5*10^5^ CFU/mL in fresh MHL and brought into the desired growth state. Stationary phase bacteria were obtained by overnight growth (16 hours). Diluted stationary phase bacteria were obtained by overnight growth (16 hours) and 1:10 dilution in fresh MHL. Exponential phase bacteria were obtained by overnight growth (16 hours), dilution to 5*10^5^ CFU/mL in fresh MHL and 3 hours of growth (Figure 2).

### Construction of knockout mutants

#### a. Vector construction

The first and last 500 bp regions of the gene (*lyta* or *spxb*) were amplified from *S. pneumoniae* D39 chromosomal DNA by PCR using Q5 High-Fidelity DNA polymerase (New England Biolabs). The kanamycin cassette was amplified from pSt-K and the streptomycin cassette from pGMC5-SM-RFP-PFurA-GFP-streptomycin. The PCR primers contained overhang sequences with the antibiotic resistance marker (kanamycin cassette for *lyta* and streptomycin cassette for *spxb*) and the vector, pGEM-T Easy vector (Promega) (*supplementary data table 6*). The first and last 500 bp of the gene and the antibiotic cassette were then introduced in the pGEM-T Easy vector using HiFi DNA assembly (New England Biolabs) resulting in plasmids pLytA and pSpxB (Figure 9) and were used to transform chemocompetent *E. coli* DH5-α. The resultant plasmid was verified by PCR and sequencing and used to transform *S. pneumoniae* D39.

**Figure 9:**
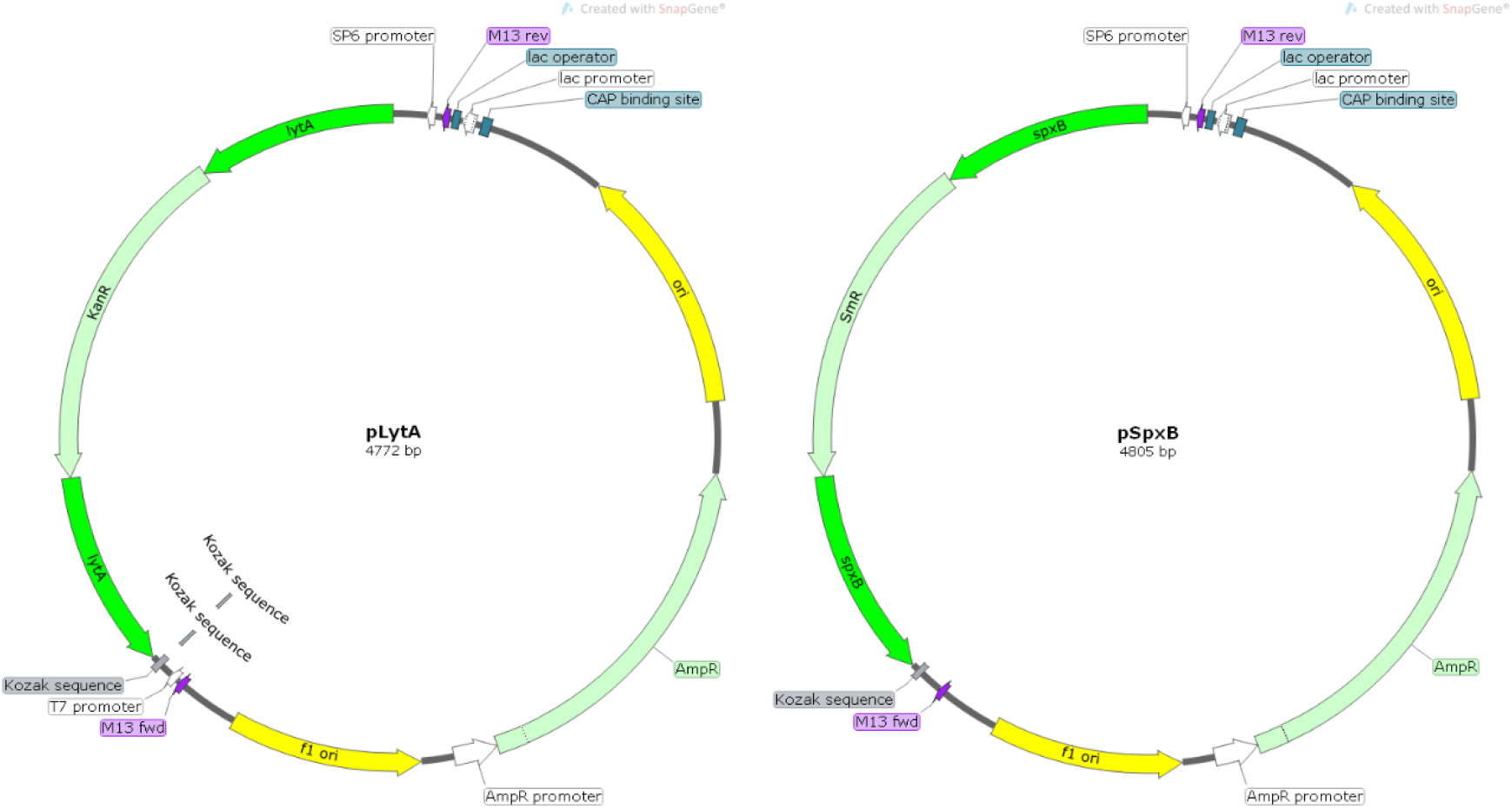
Schematic overview of the constructed plasmids to generate knockout mutants of *lyta* (pLytA) and *spxb* (pSpxB) in *S. pneumoniae* D39. The plasmid contains the first and last 500 bp of the gene (*lyta* or *spxb*) disrupted by an antibiotic resistance marker (kanamycin cassette for *lyta* and streptomycin cassette for *spxb*).

#### b. Transformation

Precompetent *S. pneumoniae* cells ware obtained by growing *S. pneumoniae* in THY to 3*10^8^ CFU/mL from a starting concentration of 1*10^6^ CFU/mL. Then, the bacterial suspension was diluted 1:100 in competence medium (THY supplemented with 0.2% bovine serum albumin and 0.01% CaCl2), 10% glycerol (Sigma-Aldrich) was added and bacteria were stored at -80°C. For transformation, precompetent *S. pneumoniae* were thawed, competence stimulating peptide 1 (CSP-1) was added (2.5 µg/mL) and competence was induced by incubation at 37°C in a water bath. After 20 minutes, 200 ng plasmid DNA was added and bacteria were incubated for an additional 60 minutes at 30°C and transferred to 37°C for 90 minutes before plating on BA containing 400 µg/mL kanamycin (*lyta*) (Sigma-Aldrich) or 200 µg/mL streptomycin (*spxb*) (Sigma-Aldrich). Resistant colonies were selected and the mutation was confirmed by sequencing. Knockdown of *lyta* and *spxb* was confirmed by qPCR (*supplementary data figure 4*).

### Antibiotic susceptibility

The minimal inhibitory concentration (MIC) of standard antibiotics was determined using a resazurin assay as described previously (49). Used antibiotics were amoxicillin (Sigma-Aldrich, beta-lactam antibiotic), cefuroxime (Sigma-Aldrich, beta-lactam antibiotic), moxifloxacin (Sigma-Aldrich, fluoroquinolone) and vancomycin (Sigma-Aldrich, glycopeptide), four clinically relevant antibiotics that are either commonly used to treat *S. pneumoniae* infections (amoxicillin, cefuroxime and moxifloxacin) or as last resort, mostly in a hospital environment (vancomycin) (80–82). Briefly, a 1:2 serial dilution of the antibiotic was made in triplicates in MHL in a 96-well plate with a final volume of 100 µL. Then, 100 µL of a bacterial suspension was added to each well, except to negative control wells, to a final concentration of 5*10^5^ CFU/mL in 200 µL. Positive control wells contained 200 µL bacterial suspension (5*10^5^ CFU/mL) without antibiotics and negative control wells contained 200 µL MHL without antibiotic or bacteria. Plates were incubated at 37°C and 5% CO_2_ for 20 hours before 20 µL of a 0.005% resazurin solution was added. Plates were further incubated for 90 minutes and fluorescence was measured (λem = 590 nm, λex = 520 nm) using a spectrophotometer (Promega Discover).

### Dose-dependent and time-kill kill curves

To obtain dose-dependent kill curves, *S. pneumoniae* was treated during 5 hours in the stationary or diluted stationary growth phase with three different antibiotic concentrations (20x, 100x and 200x MIC, respectively 0,14; 0,70 and 1,40 µg/mL for amoxicillin, 0,54; 2,7 and 5,4 µg/mL for cefuroxime, 5,8; 29 and 58 µg/mL for moxifloxacin and 8,8; 44 and 88 µg/mL for vancomycin). After 5 hours, bacterial suspensions were centrifuged, resuspended in PBS to wash away antibiotics and enumerated by VPC. Colonies were counted after minimum 48 hours of incubation. To obtain time-kill curves, *S. pneumoniae* was treated in the diluted stationary or exponential growth state with a fixed antibiotic concentration (100-fold the MIC). Bacterial suspensions were incubated during 8 or 24 hours. At specified time points, bacterial suspensions were enumerated using VPC after centrifugation and resuspension in PBS to wash away the antibiotics. Colonies were counted after minimum 48 hours of incubation.

### Heritability assay

For each condition (growth phase x antibiotic), 5 clones from the initial time-kill experiment were isolated from the blood agar plate from the second killing phase (after 6 hours of treatment for the diluted stationary phase and after 18 hours of treatment for the exponential phase), regrown in fresh MHL without antibiotic and stored at -80°C. These bacterial clones were subjected to the same protocol as the initial time-kill assay. For one of these clones per condition, a time-kill curve was obtained and the MIC value was determined. For the other 4 clones, a fixed time point (6 hours of treatment) was chosen to determine survival.

### Screening of lab strains and clinical isolates for persistence

A screening model was set-up for clinical isolates based on the protocol used for strain D39. Briefly, *S. pneumoniae* was brought into the right growth phase (diluted stationary or exponential growth phase) followed by treatment with 100-fold the MIC of amoxicillin or vancomycin. After 8 hours, bacterial suspensions were enumerated using VPC after centrifugation and resuspension in PBS to wash away the antibiotics and survival fractions were calculated.

### Data analysis and statistics

Student’s t-test, one-way ANOVA or two-way ANOVA were used to compare continuous variables (MIC values, survival fractions and time-kill curves) in Graphpad Prism Version 9. A difference between two groups was considered statistically significant when P < 0.05. The R packages *nls*.*multstart, broom* and *purrr* were used to analyze the time-kill curves mathematically by comparing two models of killing, a uniphasic model with a single killing rate and a biphasic model with two killing rates. The non-linear fixed-effect model used the Log10-transformed fraction of surviving cells. The biphasic model was based on the equation Log(Y)=log((N-P_0_)^(-kn*t)^+P_0_^(-kp*t)^) and the monophasic model on Log(Y)=log((N)^(-kn*t)^) with *Y* survival fraction, *t* treatment time (in hours), *P_0_* persister fraction at t = 0 and *k*_*n*_ and *k*_*p*_ the killing rate of normal and persister cells (per hour). Curves were considered biphasic if biphasic fitting was better than uniphasic fitting according to ANOVA (F-test), the Akaikes Information Criteria (AICs), the Bayesian information Criterion (BIC) and the Log-Likelihood (LogLik). The R package *ggcorrplot* was used to execute correlation analyses.

## ACKNOWLEDGMENTS

This research was funded by the Research Foundation-Flanders (FWO), grant numbers 12O1917N, V428917N and 1513120N; B.V.d.B. and N.G. were funded by the FWO (postdoctoral fellowship 12O1922N and predoctoral fellowship 11E4721N, respectively).

## REFERENCES

1. Subramanian K, Henriques-Normark B, Normark S. 2019. Emerging concepts in the pathogenesis of the Streptococcus pneumoniae: From nasopharyngeal colonizer to intracellular pathogen. Cell Microbiol 21.

2. Troeger C, Blacker B, Khalil IA, Rao PC, Cao J, Zimsen SRM, Albertson SB, Deshpande A, Farag T, Abebe Z, Adetifa IMO, Adhikari TB, Akibu M, Al Lami FH, Al-Eyadhy A, Alvis-Guzman N, Amare AT, Amoako YA, Antonio CAT, Aremu O, Asfaw ET, Asgedom SW, Atey TM, Attia EF, Avokpaho Efga, Ayele HT, Ayuk TB, Balakrishnan K, Barac A, Bassat Q, Behzadifar M, Behzadifar M, Bhaumik S, Bhutta ZA, Bijani A, Brauer M, Brown A, Camargos PAM, Castañeda-Orjuela Ca, Colombara D, Conti S, Dadi AF, Dandona L, Dandona R, Do HP, Dubljanin E, Edessa D, Elkout H, Endries AY, Fijabi DO, Foreman KJ, Forouzanfar MH, Fullman N, Garcia-Basteiro AL, Gessner BD, Gething PW, Gupta R, Gupta T, Hailu GB, Hassen HY, Hedayati MT, Heidari M, Hibstu DT, Horita N, Ilesanmi OS, Jakovljevic MB, Jamal AA, Kahsay A, Kasaeian A, Kassa DH, Khader YS, Khan EA, Khan MN, Khang YH, Kim YJ, Kissoon N, Knibbs LD, Kochhar S, Koul PA, Kumar GA, Lodha R, Magdy Abd El Razek H, Malta DC, Mathew JL, Mengistu DT, Mezgebe HB, Mohammad KA, Mohammed MA, Momeniha F, Murthy S, Nguyen CT, Nielsen KR, Ningrum DNA, Nirayo YL, Oren E, Ortiz Jr, PA M, Postma MJ, Qorbani M, Quansah R, Rai RK, Rana SM, Ranabhat CL, Ray SE, Rezai MS, Ruhago GM, Safiri S, Salomon JA, Sartorius B, Savic M, Sawhney M, She J, Sheikh A, Shiferaw MS, Shigematsu M, Singh JA, Somayaji R, Stanaway JD, Sufiyan MB, Taffere GR, Temsah MH, Thompson MJ, Tobe-Gai R, Topor-Madry R, Tran BX, Tran TT, Tuem KB, Ukwaja KN, Vollset SE, Walson JL, Weldegebreal F, Werdecker A, West TE, Yonemoto N, Zaki MES, Zhou L, Zodpey S, Vos T, Naghavi M, Lim SS, Mokdad AH, Murray CJL, Hay SI, Reiner RC. 2018. Estimates of the global, regional, and national morbidity, mortality, and aetiologies of lower respiratory infections in 195 countries, 1990-2016: a systematic analysis for the Global Burden of Disease Study 2016. Lancet Infect Dis 18:1191–1210.

3. Centers for Disease Control U. 2019. Antibiotic Resistance Threats in the United States, 2019.

4. Balaban NQ, Helaine S, Lewis K, Ackermann M, Aldridge B, Andersson DI, Brynildsen MP, Bumann D, Camilli A, Collins JJ, Dehio C, Fortune S, Ghigo JM, Hardt WD, Harms A, Heinemann M, Hung DT, Jenal U, Levin BR, Michiels J, Storz G, Tan MW, Tenson T, Van Melderen L, Zinkernagel A. 2019. Definitions and guidelines for research on antibiotic persistence. Nat Rev Microbiol 17:441–448.

5. Gollan B, Grabe G, Michaux C, Helaine S. 2019. Bacterial Persisters and Infection: Past, Present, and Progressing. Annu Rev Microbiol 73:359–385.

6. Windels EM, Michiels JE, Fauvart M, Wenseleers T, Bram Van Den Bergh, Michiels J. 2019. Bacterial persistence promotes the evolution of antibiotic resistance by increasing survival and mutation rates. ISME J 13:1239–1251.

7. Fauvart M, de Groote VN, Michiels J. 2011. Role of persister cells in chronic infections: clinical relevance and perspectives on anti-persister therapies. J Med Microbiol 60:699–709.

8. Santi I, Manfredi P, Maffei E, Egli A, Jenal U. 2021. Evolution of Antibiotic Tolerance Shapes Resistance Development in Chronic Pseudomonas aeruginosa Infections. MBio 12:1–17.

9. Joshi H, Kandari D, Bhatnagar R. 2021. Insights into the molecular determinants involved in Mycobacterium tuberculosis persistence and their therapeutic implications. Virulence 12:2721–2749.

10. Bakkeren E, Huisman JS, Fattinger SA, Hausmann A, Furter M, Egli A, Slack E, Sellin ME, Bonhoeffer S, Regoes RR, Diard M, Hardt WD. 2019. Salmonella persisters promote the spread of antibiotic resistance plasmids in the gut. Nature 573:276–280.

11. Kawano A, Yamasaki R, Sakakura T, Takatsuji Y, Haruyama T, Yoshioka Y, Ariyoshi W. 2020. Reactive Oxygen Species Penetrate Persister Cell Membranes of Escherichia coli for Effective Cell Killing. Front Cell Infect Microbiol 10.

12. Novak R, Henriques B, Charpentier E, Normark S, Tuomanen E. 1999. Emergence of vancomycin tolerance in Streptococcus pneumoniae. Nature 399:590–593.

13. Tomasz A, Albino A, Zanati E. 1970. Multiple antibiotic resistance in a bacterium with suppressed autolytic system. Nature 227:138–140.

14. Liu X, Li JW, Feng Z, Luo Y, Veening JW, Zhang JR. 2017. Transcriptional Repressor PtvR Regulates Phenotypic Tolerance to Vancomycin in Streptococcus pneumoniae. J Bacteriol 199.

15. Normark BH, Normark S. 2002. Antibiotic tolerance in pneumococci. Clin Microbiol Infect 8:613–622.

16. Charpentier E, Tuomanen E. 2000. Mechanisms of antibiotic resistance and tolerance in Streptococcus pneumoniae. Microbes Infect 2:1855–1864.

17. Jung SH, Ryu CM, Kim JS. 2019. Bacterial persistence: Fundamentals and clinical importance. J Microbiol 57:829–835.

18. van den Bergh B, Fauvart M, Michiels J. 2017. Formation, physiology, ecology, evolution and clinical importance of bacterial persisters. FEMS Microbiol Rev 41:219–251.

19. Oliver L, Lalier L, Salaud C, Heymann D, Cartron PF, Vallette FM. 2020. Drug resistance in glioblastoma: are persisters the key to therapy? Cancer drug Resist (Alhambra, Calif) 3:287– 301.

20. Wuyts J, Van Dijck P, Holtappels M. 2018. Fungal persister cells: The basis for recalcitrant infections? PLoS Pathog 14.

21. Jung SH, Ryu CM, Kim JS. 2019. Bacterial persistence: Fundamentals and clinical importance. J Microbiol 57:829–835.

22. Zheng EJ, Andrews IW, Grote AT, Manson AL, Alcantar MA, Earl AM, Collins JJ. 2022. Modulating the evolutionary trajectory of tolerance using antibiotics with different metabolic dependencies. Nat Commun 13.

23. Schumacher MA, Balani P, Min J, Chinnam NB, Hansen S, Vulić M, Lewis K, Brennan RG. 2015. HipBA-promoter structures reveal the basis of heritable multidrug tolerance. Nature 524:59– 66.

24. Bartell JA, Cameron DR, Mojsoska B, Haagensen JAJ, Pressler T, Sommer LM, Lewis K, Molin S, Johansen HK. 2020. Bacterial persisters in long-term infection: Emergence and fitness in a complex host environment. PLoS Pathog 16.

25. Van den Bergh B, Schramke H, Michiels JE, Kimkes TEP, Radzikowski JL, Schimpf J, Vedelaar SR, Burschel S, Dewachter L, LonČar N, Schmidt A, Meijer T, Fauvart M, Friedrich T, Michiels J, Heinemann M. 2022. Mutations in respiratory complex I promote antibiotic persistence through alterations in intracellular acidity and protein synthesis. Nat Commun 13.

26. Fuentes DAF, Manfredi P, Jenal U, Zampieri M. 2021. Pareto optimality between growth-rate and lag-time couples metabolic noise to phenotypic heterogeneity in Escherichia coli. Nat Commun 12.

27. Sebastian J, Swaminath S, Nair RR, Jakkala K, Pradhan A, Ajitkumar P. 2017. De Novo Emergence of Genetically Resistant Mutants of Mycobacterium tuberculosis from the Persistence Phase Cells Formed against Antituberculosis Drugs In Vitro. Antimicrob Agents Chemother 61.

28. Levin-Reisman I, Ronin I, Gefen O, Braniss I, Shoresh N, Balaban NQ. 2017. Antibiotic tolerance facilitates the evolution of resistance. Science 355:826–830.

29. Levin-Reisman I, Brauner A, Ronin I, Balaban NQ. 2019. Epistasis between antibiotic tolerance, persistence, and resistance mutations. Proc Natl Acad Sci U S A 116:14734–14739.

30. Huo W, Busch LM, Hernandez-Bird J, Hamami E, Marshall CW, Geisinger E, Cooper VS, van Opijnen T, Rosch JW, Isberg RR. 2022. Immunosuppression broadens evolutionary pathways to drug resistance and treatment failure during Acinetobacter baumannii pneumonia in mice. Nat Microbiol 7:796–809.

31. Restrepo A V., Salazar BE, Agudelo M, Rodriguez CA, Zuluaga AF, Vesga O. 2005. Optimization of culture conditions to obtain maximal growth of penicillin-resistant Streptococcus pneumoniae. BMC Microbiol 5.

32. Regev-Yochay G, Trzcinski K, Thompson CM, Lipsitch M, Malley R. 2007. SpxB is a suicide gene of Streptococcus pneumoniae and confers a selective advantage in an in vivo competitive colonization model. J Bacteriol 189:6532–6539.

33. Mellroth P, Daniels R, Eberhardt A, Rönnlund D, Blom H, Widengren J, Normark S, Henriques-Normark B. 2012. LytA, Major Autolysin of Streptococcus pneumoniae, Requires Access to Nascent Peptidoglycan https://doi.org/10.1074/jbc.M111.318584.

34. Bryant JC, Dabbs RC, Oswalt KL, Brown LR, Rosch JW, Seo KS, Donaldson JR, McDaniel LS, Thornton JA. 2016. Pyruvate oxidase of Streptococcus pneumoniae contributes to pneumolysin release. BMC Microbiol 16:1–12.

35. Regev-Yochay G, Trzciński K, Thompson CM, Malley R, Lipsitch M. 2006. Interference between Streptococcus pneumoniae and Staphylococcus aureus: In vitro hydrogen peroxide-mediated killing by Streptococcus pneumoniae. J Bacteriol 188:4996–5001.

36. Lisher JP, Tsui H-CT, Ramos-Montañez S, Hentchel KL, Martin JE, Trinidad JC, Winkler ME, Giedroc DP. 2017. Biological and Chemical Adaptation to Endogenous Hydrogen Peroxide Production in Streptococcus pneumoniae D39. mSphere 2.

37. Martner A, Skovbjerg S, Paton JC, Wold AE. 2009. Streptococcus pneumoniae autolysis prevents phagocytosis and production of phagocyte-activating cytokines. Infect Immun 77:3826–3837.

38. Giudicelli S, Tomasz A. 1984. Attachment of Pneumococcal Autolysin to Wall Teichoic Acids, an Essential Step in Enzymatic Wall Degradation. J Bacteriol 158:1188–1190.

39. Herta T, Bhattacharyya A, Bollensdorf C, Kabus C, García P, Suttorp N, Hippenstiel S, Zahlten J. 2018. DNA-release by Streptococcus pneumoniae autolysin LytA induced Krueppel-like factor 4 expression in macrophages. Sci Rep 8.

40. Verhagen LM, de Groot R. 2015. Recurrent, protracted and persistent lower respiratory tract infection: A neglected clinical entity. J Infect 71:S106–S111.

41. Hare KM, Leach AJ, Smith-Vaughan HC, Chang AB, Grimwood K. 2017. Streptococcus pneumoniae and chronic endobronchial infections in childhood. Pediatr Pulmonol 52:1532–1545.

42. Inchingolo R, Pierandrei C, Montemurro G, Smargiassi A, Lohmeyer FM, Rizzi A. 2021. Antimicrobial Resistance in Common Respiratory Pathogens of Chronic Bronchiectasis Patients: A Literature Review. Antibiot (Basel, Switzerland) 10.

43. Lewis K. 2010. Persister cells. Annu Rev Microbiol 64:357–372.

44. Hoa M, Tomovic S, Nistico L, Hall-Stoodley L, Stoodley P, Sachdeva L, Berk R, Coticchia JM. 2009. Identification of adenoid biofilms with middle ear pathogens in otitis-prone children utilizing SEM and FISH. Int J Pediatr Otorhinolaryngol 73:1242–1248.

45. Hall-Stoodley L, Hu FZ, Gieseke A, Nistico L, Nguyen D, Hayes J, Forbes M, Greenberg DP, Dice B, Burrows A, Wackym PA, Stoodley P, Post JC, Ehrlich GD, Kerschner JE. 2006. Direct detection of bacterial biofilms on the middle-ear mucosa of children with chronic otitis media. JAMA 296:202–211.

46. Nistico L, Kreft R, Gieseke A, Coticchia JM, Burrows A, Khampang P, Liu Y, Kerschner JE, Post JC, Lonergan S, Sampath R, Hu FZ, Ehrlich GD, Stoodley P, Hall-Stoodley L. 2011. Adenoid reservoir for pathogenic biofilm bacteria. J Clin Microbiol 49:1411–1420.

47. Lewis K. 2008. Multidrug tolerance of biofilms and persister cells. Curr Top Microbiol Immunol 322:107–131.

48. Suárez N, Texeira E. 2019. Optimal Conditions for Streptococcus pneumoniae Culture: In Solid and Liquid Media. Methods Mol Biol 1968:3–10.

49. Cools F, Torfs E, Vanhoutte B, de Macedo MB, Bonofiglio L, Mollerach M, Maes L, Caljon G, Delputte P, Cappoen D, Cos P. 2018. Streptococcus pneumoniae galU gene mutation has a direct effect on biofilm growth, adherence and phagocytosis in vitro and pathogenicity in vivo. Pathog Dis 76.

50. Luidalepp H, Jõers A, Kaldalu N, Tenson T. 2011. Age of inoculum strongly influences persister frequency and can mask effects of mutations implicated in altered persistence. J Bacteriol 193:3598–3605.

51. Keren I, Kaldalu N, Spoering A, Wang Y, Lewis K. 2004. Persister cells and tolerance to antimicrobials. FEMS Microbiol Lett 230:13–18.

52. Grant SS, Hung DT. 2013. Persistent bacterial infections, antibiotic tolerance, and the oxidative stress response. Virulence 4:273–283.

53. Michiels JE, Van Den Bergh B, Verstraeten N, Fauvart M, Michiels J. 2016. In Vitro Emergence of High Persistence upon Periodic Aminoglycoside Challenge in the ESKAPE Pathogens. Antimicrob Agents Chemother 60:4630–4637.

54. Ma D, Mandell JB, Donegan NP, Cheung AL, Ma W, Rothenberger S, Shanks RMQ, Richardson AR, Urish KL. 2019. The Toxin-Antitoxin MazEF Drives Staphylococcus aureus Biofilm Formation, Antibiotic Tolerance, and Chronic Infection. MBio 10.

55. Savijoki K, Miettinen I, Nyman TA, Kortesoja M, Hanski L, Varmanen P, Fallarero A. 2020. Growth Mode and Physiological State of Cells Prior to Biofilm Formation Affect Immune Evasion and Persistence of Staphylococcus aureus. Microorganisms 8.

56. Suppiger S, Astasov-Frauenhoffer M, Schweizer I, Waltimo T, Kulik EM. 2020. Tolerance and Persister Formation in Oral Streptococci. Antibiot (Basel, Switzerland) 9.

57. Leung V, Lévesque CM. 2012. A stress-inducible quorum-sensing peptide mediates the formation of persister cells with noninherited multidrug tolerance. J Bacteriol 194:2265–2274.

58. Jiang Y-L, Qiu W, Zhou X-D, Li H, Lu J-Z, Xu HH, Peng X, Li M-Y, Feng M-Y, Cheng L, Ren B. 2017. Quaternary ammonium-induced multidrug tolerant Streptococcus mutans persisters elevate cariogenic virulence in vitro. Int J Oral Sci 9:e7–e7.

59. Wang S, Zhou C, Ren B, Li X, Weir MD, Masri RM, Oates TW, Cheng L, Xu HKH. 2017. Formation of persisters in Streptococcus mutans biofilms induced by antibacterial dental monomer. J Mater Sci Mater Med 28:178.

60. Dufour D, Mankovskaia A, Chan Y, Motavaze K, Gong S-G, Lévesque CM. 2018. A tripartite toxin-antitoxin module induced by quorum sensing is associated with the persistence phenotype in Streptococcus mutans. Mol Oral Microbiol 33:420–429.

61. Willenborg J, Willms D, Bertram R, Goethe R, Valentin-Weigand P. 2014. Characterization of multi-drug tolerant persister cells in Streptococcus suis. BMC Microbiol 14:120.

62. Soriano F, Greenwood D. 1979. Action and interaction of penicillin and gentamicin on enterococci. J Clin Pathol 32:1174–1179.

63. Wood DN, Chaussee MA, Chaussee MS, Buttaro BA. 2005. Persistence of Streptococcus pyogenes in stationary-phase cultures. J Bacteriol 187:3319–3328.

64. Martini CL, Coronado AZ, Melo MCN, Gobbi CN, Lopez ÚS, de Mattos MC, Amorim TT, Botelho AMN, Vasconcelos ATR, Almeida LGP, Planet PJ, Zingali RB, Figueiredo AMS, Ferreira-Carvalho BT. 2021. Cellular Growth Arrest and Efflux Pumps Are Associated With Antibiotic Persisters in Streptococcus pyogenes Induced in Biofilm-Like Environments. Front Microbiol 12.

65. Thornton RB, Rigby PJ, Wiertsema SP, Filion P, Langlands J, Coates HL, Vijayasekaran S, Keil AD, Richmond PC. 2011. Multi-species bacterial biofilm and intracellular infection in otitis media. BMC Pediatr 11.

66. Verstraete L, Van den Bergh B, Verstraeten N, Michiels J. 2022. Ecology and evolution of antibiotic persistence. Trends Microbiol 30:466–479.

67. Pham TDM, Ziora ZM, Blaskovich MAT. 2019. Quinolone antibiotics. Medchemcomm 10:1719–1739.

68. Mok WWK, Brynildsen MP. 2018. Timing of DNA damage responses impacts persistence to fluoroquinolones. Proc Natl Acad Sci U S A 115:E6301–E6309.

69. Stewart B, Rozen DE. 2012. Genetic variation for antibiotic persistence in Escherichia coli. Evolution 66:933–939.

70. Hofsteenge N, Van Nimwegen E, Silander OK. 2013. Quantitative analysis of persister fractions suggests different mechanisms of formation among environmental isolates of E. coli. BMC Microbiol 13.

71. Barth VC, Rodrigues BA, Bonatto GD, Gallo SW, Pagnussatti VE, Ferreira CAS, De Oliveira SD. 2013. Heterogeneous persister cells formation in Acinetobacter baumannii. PLoS One 8.

72. Kaldalu N, Hauryliuk V, Tenson T. 2016. Persisters-as elusive as ever. Appl Microbiol Biotechnol 100:6545–6553.

73. Orman MA, Brynildsen MP. 2013. Dormancy is not necessary or sufficient for bacterial persistence. Antimicrob Agents Chemother 57:3230–3239.

74. Hill PWS, Moldoveanu AL, Sargen M, Ronneau S, Glegola-Madejska I, Beetham C, Fisher RA, Helaine S. 2021. The vulnerable versatility of Salmonella antibiotic persisters during infection. Cell Host Microbe 29:1757-1773.e10.

75. Shan Y, Gandt AB, Rowe SE, Deisinger JP, Conlon BP, Lewis K. 2017. ATP-Dependent Persister Formation in Escherichia coli. MBio 8.

76. Conlon BP, Rowe SE, Gandt AB, Nuxoll AS, Donegan NP, Zalis EA, Clair G, Adkins JN, Cheung AL, Lewis K. 2016. Persister formation in Staphylococcus aureus is associated with ATP depletion. Nat Microbiol 1.

77. Stapels DAC, Hill PWS, Westermann AJ, Fisher RA, Thurston TL, Saliba AE, Blommestein I, Vogel J, Helaine S. 2018. Salmonella persisters undermine host immune defenses during antibiotic treatment. Science 362:1156–1160.

78. Peyrusson F, Varet H, Nguyen TK, Legendre R, Sismeiro O, Coppée JY, Wolz C, Tenson T, Van Bambeke F. 2020. Intracellular Staphylococcus aureus persisters upon antibiotic exposure. Nat Commun 11.

79. Goneau LW, Yeoh NS, MacDonald KW, Cadieux PA, Burton JP, Razvi H, Reid G. 2014. Selective target inactivation rather than global metabolic dormancy causes antibiotic tolerance in uropathogens. Antimicrob Agents Chemother 58:2089–2097.

80. De Sutter A, Piessens V, Poelman P, Van Roy K. 2021. BCFI | Methodologie van de update van de gids 2019/2021. https://www.bcfi.be/nl/chapters/12?matches=BAPCOC&frag=8901316. Retrieved 25 February 2022.

81. Rodriguez CA, Atkinson R, Bitar W, Whitney CG, Edwards KM, Mitchell L, Li J, Sublett J, Li CS, Liu T, Chesney FJ, Tuomanen EI. 2004. Tolerance to vancomycin in pneumococci: detection with a molecular marker and assessment of clinical impact. J Infect Dis 190:1481–1487.

82. Corrêa R de A, Costa AN, Lundgren F, Michelin L, Figueiredo MR, Holanda M, Gomes M, Teixeira PJZ, Martins R, Silva R, Athanazio RA, da Silva RM, Pereira MC. 2018. 2018 recommendations for the management of community acquired pneumonia. J Bras Pneumol 44:405–423.

