## Supplementary data for "Antibiotic-tolerant persisters are pervasive among clinical *Streptococcus pneumoniae* isolates and show strong condition-dependence"

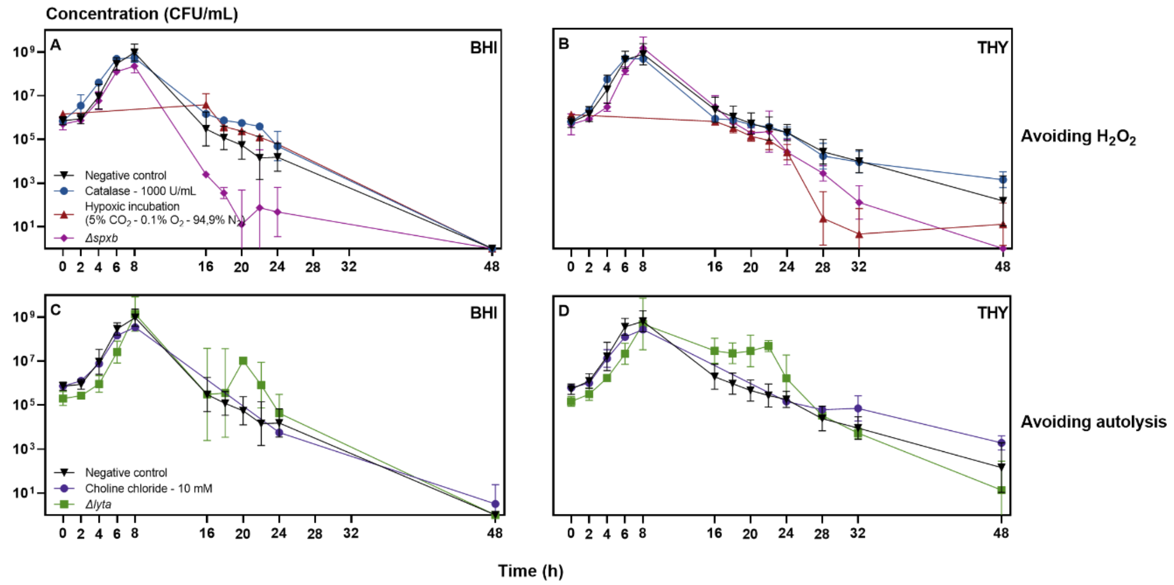


Figure 1: **The self-limiting *in vitro* nature of *S. pneumoniae* is not counteracted in BHI and THY despite the different strategies adopted to avoid $H_2O_2$ or autolysis.**
We compared the effect of different strategies to counteract the effects of pyruvate oxidase (A and B) or autolysin (C and D) in planktonic growth curves of *S. pneumoniae* D39 in BHI (Brain Heart Infusion broth, A and C) and THY (Todd-Hewitt broth supplemented with 0.5% Yeast extract, B and D). The strong reduction of viable bacteria after 8 hours of growth is still observed despite the adopted strategies. The experiments were performed in triplicates and each value is presented as the mean ± standard deviation (n = 3).

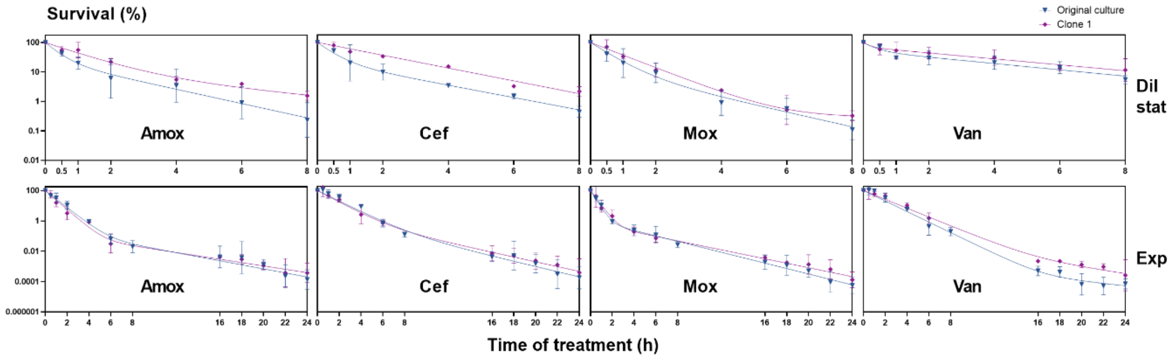


Figure 2: **The antibiotic tolerance of surviving *S. pneumoniae* cells is transient and non-deterministically inherited by daughter cells.**
Fitting of a non-linear fixed-effect model to log-transformed kill curves of amoxicillin (amox), cefuroxime (cef), moxifloxacin (mox) and vancomycin (van) against *S. pneumoniae* D39 planktonic bacteria. AB-tolerant *S. pneumoniae* D39 clones were recovered after 6 (Dil stat) or 18 (Exp) hours of treatment during the initial time-kill assay, regrown without antibiotic and preserved at -80°C. For one of these clones arising from potential persister cells, survival was determined over 8 or 24 hours of antibiotic treatment with amoxicillin (amox), cefuroxime (cef), moxifloxacin (mox) and vancomycin (van) in the diluted stationary (Dil stat) or the exponential growth phase (Exp). Killing dynamic patterns of the randomly selected clones were similar to the original culture (two-way ANOVA). The experiments were performed in triplicates and each value is presented as the mean ± standard deviation (n = 3).

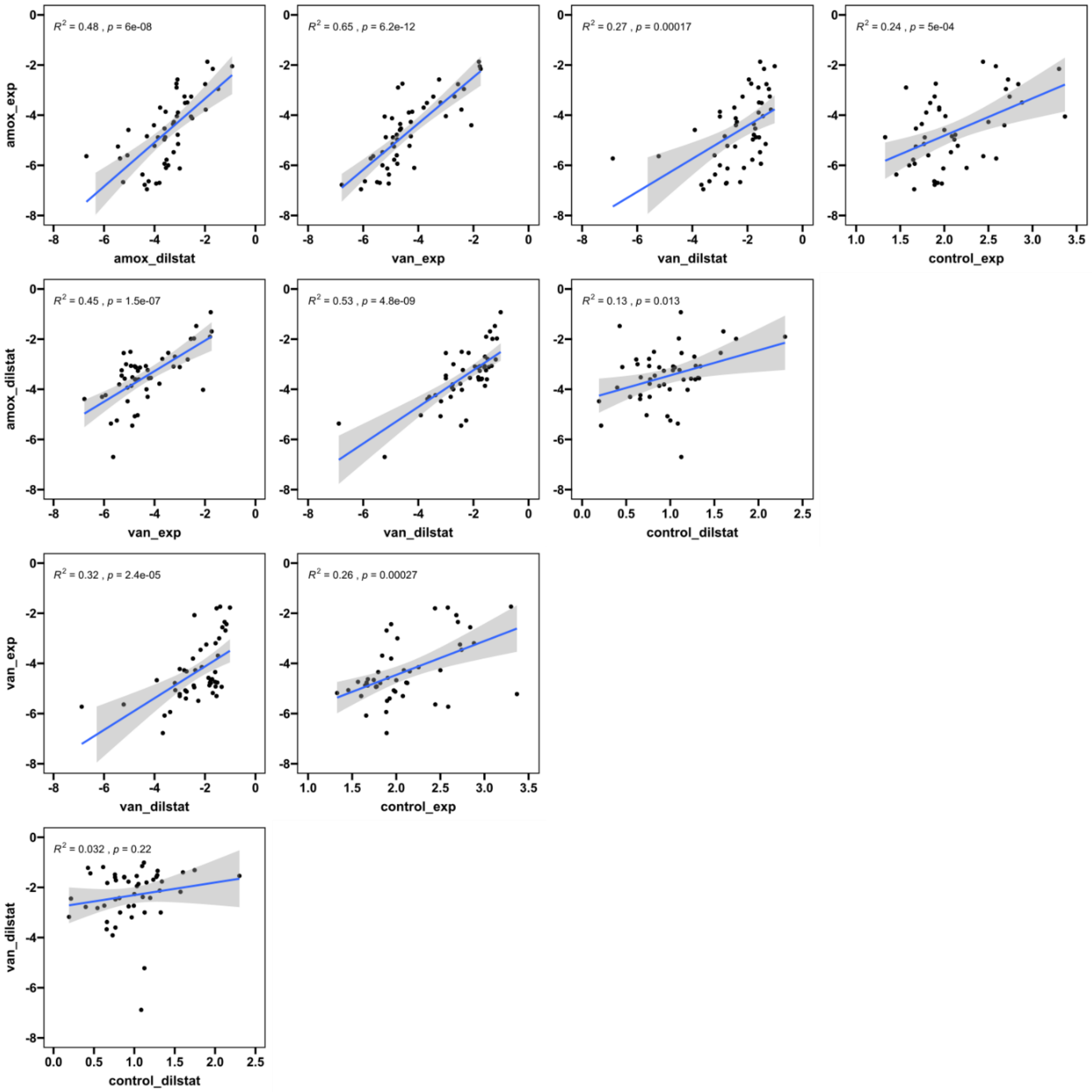


Figure 3: **Correlation analysis of survival fractions between different conditions show strong correlations between antibiotics (amoxicillin and vancomycin) and growth phases (diluted stationary and exponential growth phase).**

Individual correlations between the survival rates in 4 different conditions: treatment with amoxicillin (amox) or vancomycin (van) in the diluted stationary (dilstat) or exponential (exp) growth phase. Also the correlation with the corresponding control (exponential or diluted stationary growth) is given. Pearson correlation coefficients ($R^2$) are given for each correlation.

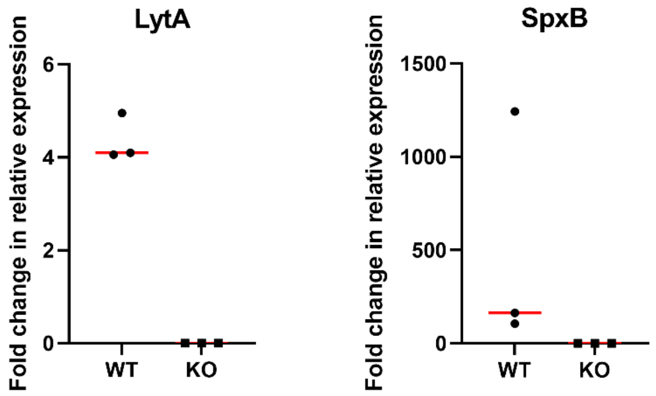


Figure 4: **The knockout mutants of the enzymes autolysin (LytA) and pyruvate oxidase (SpxB) show mRNA expression.**
The data represent the number of fold-changes in mRNA levels of knockout mutants (KO) and wild-types (WT). The mean is given as the red line with the individual datapoints as dots (n = 1x3).

Table 1: ***S. pneumoniae* D39 is susceptible to the antibiotics amoxicillin, cefuroxime, moxifloxacin and vancomycin according to the EUCAST breaking points.**
Minimum inhibitory concentration (MIC) of reference strain D39. Values represent mean ± SD (n = 3).

| MIC (µg/mL) | D39 | EUCAST breaking points | |
|---|---|---|---|
| | | Sensitive (µg/mL) | Resistant (µg/mL) |
| **Amoxicillin** | 0.007 ± 0.002 | ≤ 0.5 | ≥ 1 |
| **Cefuroxime** | 0.022 ± 0.005 | ≤ 0.25 | > 0.5 |
| **Moxifloxacin** | 0.233 ± 0.006 | ≤ 0.5 | ≥ 0.5 |
| **Vancomycin** | 0.450 ± 0.111 | ≤ 2 | ≥ 2 |

Table 2: **The biphasic model describes the time-resolved killing data better than the uniphasic model.** Mathematical analyses of the entire dataset, with a global model containing a condition-dependent structure, and of the individual conditions by comparing the fitting of two non-linear fixed-effect models (uniphasic versus biphasic model) to time-kill curves. Probability that the model is correct is determined using the Akaike's information criterion (AICc), the Bayesian information Criterion (BIC) and the Log-Likelihood (LogLik). If the AICc/BIC are lower and the LogLik is higher, the probability is higher that the model is correct. Dil stat, diluted stationary growth phase; Exp, exponential growth phase.

| | AIC | BIC | LogLik | AIC | BIC | LogLik |
|---|---|---|---|---|---|---|
| **Global model comparison (across all treatments)** | | | | | | |
| Uniphasic | 915.8 | 951.2 | -448.9 | | | |
| Biphasic | 481.4 | 579.8 | -215.7 | | | |
| **Amoxicillin** | **Exp** | | | **Dil stat** | | |
| Uniphasic | 204.1 | 208.5 | -100.1 | 36.3 | 38.9 | -16.2 |
| Biphasic | 115.7 | 124.6 | -53.9 | 33.0 | 38.2 | -12.5 |
| **Cefuroxime** | **Exp** | | | **Dil stat** | | |
| Uniphasic | 146.2 | 150.7 | -71.1 | 28.9 | 31.5 | -12.4 |
| Biphasic | 104.6 | 113.5 | -48.3 | 28.8 | 34.0 | -10.4 |
| **Moxifloxacin** | **Exp** | | | **Dil stat** | | |
| Uniphasic | 219.0 | 223.4 | -107.5 | 25.0 | 27.6 | -10.5 |
| Biphasic | 88.8 | 97.7 | -40.4 | 18.3 | 23.5 | -5.2 |
| **Vancomycin** | **Exp** | | | **Dil stat** | | |
| Uniphasic | 118.4 | 122.9 | -57.2 | -7.2 | -4.6 | 5.6 |
| Biphasic | 68.7 | 77.6 | -30.4 | -8.2 | -3.0 | 8.1 |

Table 3: **Mathematical analysis of the fitting of a biphasic non-linear fixed-effect model to kill curves of amoxicillin (amox), cefuroxime (cef), moxifloxacin (mox) and vancomycin (van) against *S. pneumoniae* D39.** 95% confidence intervals of the parameters are given between brackets. $P_0$, persister fraction at the start of treatment; $K_n$ , killing rate of normal cells; $K_P$, killing rate of persister cells; Exp, exponential growth phase; Dil stat, diluted stationary growth phase.

| Antibiotic | Growth phase | $P_0$ | $K_n$ | $K_p$ |
|---|---|---|---|---|
| **Amox** | Exp | 0.0012 (-0.0098 – 0.0034) | 1.3240 (1.1281 – 1.5200) | 0.2628 (0.1703 – 0.3553) |
| | Dil stat | 0.2431 (-0.1645 – 0.6507) | 2.5802 (-1.3650 – 6.5254) | 0.5586 (0.2708 – 0.8463) |
| **Cef** | Exp | 0.0050 (-0.0117 – 0.0216) | 0.9259 (0.7873 – 1.0644) | 0.3166 (0.1566 – 0.4765) |
| | Dil stat | 0.4650 (-0.0644 – 0.9944) | 3.7800 (-10.5042 – 18.0642) | 0.5067 (0.2981 – 0.7153) |
| **Mox** | Exp | 0.0040 (0.0014 – 0.0065) | 2.4595 (2.0477 – 2.8713) | 0.3611 (0.3187 – 0.4035) |
| | Dil stat | 0.1374 (-0.1644 – 0.4391) | 1.7510 (0.5172 – 2.9848) | 0.5756 (0.2320 – 0.9192) |
| **Van** | Exp | 0.0002 (-0.0005 – 0.0009) | 0.8854 (0.8194 – 0.9513) | 0.2510 (0.0962 – 0.4058) |
| | Dil stat | 0.6008 (0.1265 – 1.0750) | 2.0722 (-3.5989 – 7.7432) | 0.2706 (0.1353 – 0.4059) |

Table 4: **Minimum inhibitory concentration (MIC) of *S. pneumoniae* D39 before and after the initial time-kill curve experiment.** Values represent mean ± SD (n = 3). The MIC value was determined for one randomly selected surviving clone of the initial time-kill assay. MIC values before and after the initial time-kill experiment did not significantly differ (Student's T-test), except for the MIC for moxifloxacin in the diluted stationary growth phase that was significantly lower for the repeated experiment (p = 0.001).

| MIC (µg/mL) | D39 | Clone after antibiotic treatment in diluted stationary phase | Clone after antibiotic treatment in exponential phase |
|---|---|---|---|
| **Amoxicillin** | 0,007 ± 0,002 | 0,009 ± 0,004 | 0,010 ± 0,002 |
| **Cefuroxime** | 0,022 ± 0,005 | 0,031 ± 0,003 | 0,034 ± 0,018 |
| **Moxifloxacin** | 0,233 ± 0,006 | **0,071 ± 0,039** | 0,282 ± 0,075 |
| **Vancomycin** | 0,450 ± 0,111 | 0,426 ± 0,080 | 0,588 ± 0,155 |

Table 5: **Minimum inhibitory concentration (MIC) of reference strains and clinical isolates.** All strains are sensitive to amoxicillin (amox), cefuroxime (cef), moxifloxacin (mox) and vancomycin (van) according to the EUCAST breaking points, except for strain 85 that displays resistance towards cefuroxime and for CI 7 that displays minor resistance towards moxifloxacin. Values represent mean ± SD (n = 3).

| Strain | Amox | Cef | Mox | Van | Strain | Amox | Cef | Mox | Van |
|---|---|---|---|---|---|---|---|---|---|
| **D39** | 0,007 ± 0,002 | 0,022 ± 0,005 | 0,233 ± 0,006 | 0,450 ± 0,111 | **CI 3** | 0.014 ± 0.008 | 0.019 ± 0.006 | 0.073 ± 0.026 | 0.348 ± 0.101 |
| **TIGR4** | 0.006 ± 0.002 | 0.032 ± 0.013 | 0.347 ± 0.069 | 0.424 ± 0.092 | **CI 4** | 0.010 ± 0.003 | 0.018 ± 0.007 | 0.118 ± 0.006 | 0.365 ± 0.087 |
| **ATCC49619** | 0.036 ± 0.008 | 0.239 ± 0.145 | 0.242 ± 0.004 | 0.245 ± 0.002 | **CI 5** | 0.012 ± 0.004 | 0.017 ± 0.008 | 0.376 ± 0.096 | 0.535 ± 0.072 |
| **R6** | 0.011 ± 0.002 | 0.016 ± 0.007 | 0.226 ± 0.061 | 0.216 ± 0.017 | **CI 6** | 0.014 ± 0.001 | 0.015 ± 0.000 | 0.015 ± 0.000 | 0.496 ± 0.006 |
| **85** | 0.359 ± 0.105 | **5.215 ± 0.895** | 0.094 ± 0.026 | 0.301 ± 0.084 | **CI 7** | 0.010 ± 0.004 | 0.012 ± 0.003 | **0.637 ± 0.236** | 0.297 ± 0.106 |
| **88** | 0.009 ± 0.004 | 0.083 ± 0.020 | 0.146 ± 0.027 | 0.246 ± 0.024 | **CI 8** | 0.011 ± 0.004 | 0.013 ± 0.002 | 0.311 ± 0.048 | 0.567 ± 0.097 |
| **CI 1** | 0.014 ± 0.001 | 0.030 ± 0.000 | 0.459 ± 0.131 | 0.401 ± 0.033 | **CI 9** | 0.011 ± 0.002 | 0.022 ± 0.006 | 0.200 ± 0.005 | 0.517 ± 0.037 |
| **CI 2** | 0.124 ± 0.017 | 0.115 ± 0.007 | 0.084 ± 0.028 | 0.487 ± 0.007 | **CI 10** | 0.013 ± 0.003 | 0.012 ± 0.004 | 0.145 ± 0.030 | 0.644 ± 0.190 |

Table 6: **Primers used for plasmid construction and validation of the *spxb* and *lyta* knockout mutants.**

| Primer | Sequence |
|---|---|
| **Construction of pLytA** | |
| For_lyta_first500 | 5'-TCCCGTTGAATATGGCTCATCCATTTAGCAAGATATGGATAAGGGTCAAC-3' |
| Rev_lyta_first500 | 5'- TATGGTCGACCTGCAGGCGGCCGCGAATTCACTAGTGATTATGGAAATTAATGTGAGTAA ATTAAGAACAGATTTGCCTCAAGT -3' |
| For_kan | 5'- ATCCATATCTTGCTAAATGGATGAGCCATATTCAACGGGAAACG -3' |
| Rev_kan | 5'- TTCTCAATATCATGCTTAAATTAGAAAAACTCATCGAGCATCAAATGAAACT -3' |
| For_lyta_last500 | 5'- GCATGCTCCCGGCCGCCATGGCGGCCGCGGGAATTCGATTTTATTTTACTGTAATCAAGC CATCTGGCTCTACT- 3' |
| Rev_lyta_last500 | 5'- TGCTCGATGAGTTTTTCTAATTTAAGCATGATATTGAGAACGGCTTGAC -3' |
| **Construction of pSpxB** | |
| For_spxb_first500 | 5'- TATGGTCGACCTGCAGGCGGCCGCGAATTCACTAGTGATTATGACTCAAGGGAAAATTAC TGCATCTG -3' |
| Rev_ spxb_first500 | 5'- GCGATCACCGCTTCCCTCATGAAGTTTACTGGAATTTCAACAACAGCTGG -3' |
| For_strep | 5'- CGATCTGGATTGTCTTTCTTTTATTTGCCGACTACCTTGGTGATCT -3' |
| Rev_strep | 5'- TTGAAATTCCAGTAAACTTCATGAGGGAAGCGGTGATCGCC -3' |
| For_ spxb _last500 | 5'- CCAAGGTAGTCGGCAAATAAAAGAAAGACAATCCAGATCGCCAAG -3' |
| Rev_ spxb _last500 | 5'- GCATGCTCCCGGCCGCCATGGCGGCCGCGGGAATTCGATTTTATTTAATTGCGCGTGATT GCAATCCTTCTTCTTCCA -3' |
| **cPCR to check integration** | |
| For_lyta_cPCR | 5'- TGCGCTGTTCTGATTTGAAAGA -3' |
| Rev_lyta_cPCR | 5'- AAAGGAGTTTCTGGTTCTGGAT -3' |
| For_spxb_cPCR | 5'- TATGGTCGACCTGCAGGCGGCCGCGAATTCACTAGTGATTATGACTCAAGGGAAAATTAC TGCATCTG -3' |
| Rev_spxb_cPCR | 5'- GCATGCTCCCGGCCGCCATGGCGGCCGCGGGAATTCGATTTTATTTAATTGCGCGTGATT GCAATCCTTCTTCTTCCA -3' |
| **qPCR to check expression** | |
| For_lyta_qPCR | 5'- CAGATTTGCCTCAAGTCGGC -3' |
| Rev_lyta_qPCR | 5'- ATTCTGGGTCTTTCCGCCAG -3' |
| For_spxb_qPCR | 5'- TCTCCGCTCTTTGCGACAAT -3' |
| Rev_spxb_qPCR | 5'- TGTTGAATGCTCCATCACCCA -3' |
| For_gdh | 5'- GGAGACCTGGCTAAACGCAA -3' |
| Rev_gdh | 5'- GGTCTACGGGCAGTTCCAAT -3' |
